# The HCM – Linked Mutation Arg92Leu in TNNT2 Allosterically Alters the cTnC – cTnI Interface and Disrupts the PKA-mediated Regulation of Myofilament Relaxation

**DOI:** 10.1101/2023.07.18.549569

**Authors:** Melissa L. Lynn, Jesus Jimenez, Romi L. Castillo, Matthew M. Klass, Catherine Vasquez, Anthony Baldo, Cyonna Gibson, Anne M. Murphy, Jil C. Tardiff

## Abstract

**Background:** Impaired left ventricular relaxation, high filling pressures, and dysregulation of Ca^2+^ homeostasis are common findings contributing to diastolic dysfunction in hypertrophic cardiomyopathy (HCM). Studies have shown that impaired relaxation is an early observation in the sarcomere-gene-positive preclinical HCM cohort which suggests potential involvement of myofilament regulators of relaxation. Yet, a molecular level understanding of mechanism(s) at the level of the myofilament is lacking. We hypothesized that mutation-specific, allosterically mediated, changes to the cardiac troponin C-cardiac troponin I (cTnC-cTnI) interface can account for the development of early-onset diastolic dysfunction via decreased PKA accessibility to cTnI.

**Methods:** HCM mutations R92L-cTnT (Arg92Leu) and Δ160E-cTnT (Glu160 deletion) were studied *in vivo*, *in vitro,* and *in silico* via 2D echocardiography, western blotting, *ex vivo* hemodynamics, stopped-flow kinetics, time resolved fluorescence resonance energy transfer (TR-FRET), and molecular dynamics simulations.

**Results:** The HCM-causative mutations R92L-cTnT and Δ160E-cTnT result in different time-of-onset of diastolic dysfunction. R92L-cTnT demonstrated early-onset diastolic dysfunction accompanied by a localized decrease in phosphorylation of cTnI. Constitutive phosphorylation of cTnI (cTnI-D_23_D_24_) was sufficient to recover diastolic function to Non-Tg levels only for R92L-cTnT. Mutation-specific changes in Ca^2+^ dissociation rates associated with R92L-cTnT reconstituted with cTnI-D_23_D_24_ led us to investigate potential involvement of structural changes in the cTnC-cTnI interface as an explanation for these observations. We probed the interface via TR-FRET revealing a repositioning of the N-terminus of cTnI, closer to cTnC, and concomitant decreases in distance distributions at sites flanking the PKA consensus sequence. Implementing TR-FRET distances as constraints into our atomistic model identified additional electrostatic interactions at the consensus sequence.

**Conclusion:** These data indicate that the early diastolic dysfunction observed in a subset of HCM is likely attributable to structural changes at the cTnC-cTnI interface that impair accessibility of PKA thereby blunting β-adrenergic responsiveness and identifying a potential molecular target for therapeutic intervention.

## INTRODUCTION

Hypertrophic cardiomyopathy (HCM) is a complex, progressive disorder originally defined as left ventricular (LV) hypertrophy in the absence of other identifiable etiology ^1^. It is the most common genetic cardiac disorder, with an estimated frequency of ∼ 1/200 - 1/500 individuals worldwide ^2^. In the early 1990’s, genetic linkage studies identified mutations in the protein components of the cardiac sarcomere as the primary molecular cause of HCM ^3,4^. Subsequently, the availability of genetic testing coupled to advanced imaging techniques facilitated the first longitudinal clinical studies of genotyped cohorts, transforming our understanding of the complex, often decades-long progressive ventricular remodeling that significantly impacts management and outcome ^5,6^. These important advances, however, have raised new considerations. In particular, cascade genetic testing of first-degree relatives (now a Class I indication in HCM) has identified a preclinical, phenotypically “negative” patient cohort ^7,8^. As treatments for HCM are largely palliative and reserved for patients with symptoms that often manifest when irreversible cardiac remodeling has occurred, the ability to alter the pathogenic trajectory remains limited. Thus, the ability to identify HCM patients in the earliest stages of disease presents a unique opportunity to intervene and mitigate outcomes.

Challenges remain regarding implementation of this approach. For example, the choice of clinical target, the characterization of the primary molecular mechanism and the timing of treatment are all central considerations. The recent development and FDA approval of Mavacamten, a first-in-class targeted myosin inhibitor provides a framework for addressing these questions. Mavacamten treats symptomatic obstructive HCM by inhibiting myosin ATPase and modulating the hemodynamic effects of the LV outflow tract gradient by decreasing cardiac hypercontractility ^9,10^. By definition, it is targeted at patients in later stages of the cardiac remodeling when significant and dynamic symptoms are established. Another common manifestation of HCM is impaired LV relaxation leading to increased LV filling pressures and diastolic dysfunction. Diastolic dysfunction is a major component of the most prominent symptom in HCM, dyspnea on exertion. The etiology of impaired relaxation in HCM is multifactorial, including microvascular disease, increased LV mass, fibrosis, impaired energetics, and dysregulation of myocellular Ca^2+^ homeostasis; alone and in combination ^8^. Many of these manifestations are later in onset and represent secondary (post-myofilament) myocellular or non-myocyte-based processes that directly impact cardiac remodeling and reduce compliance. It has long been noted that diastolic dysfunction in HCM is often out-of-proportion to the degree of LV hypertrophy, this “uncoupling” supports a primary cellular mechanism ^11^. Recent studies have shown that impaired relaxation is one of the earliest observations in HCM, including preclinical sarcomeric-gene-positive cohorts, suggesting a potential proximal relationship to the myofilament and eventually cardiac diastolic dysfunction ^12-14^. In the aggregate, these observations support the premise that a component of early-onset diastolic dysfunction at the myocellular level resides in the cardiac myofilament and thus, like cardiac hypercontractility, may be directly targetable.

While the majority of sarcomeric HCM has been linked to mutations in myosin binding protein C (MYBPC3) and beta myosin heavy chain (MYH7), mutations in cardiac thin filament (CTF) proteins often result in complex, difficult to manage non-obstructive forms of HCM and, occasionally RCM ^15,16^. The CTF is the essential molecular regulator of the actomyosin crossbridge cycle. It is comprised of five discrete proteins: cTnC, cTnI, cTnT, actin and tropomyosin that co-evolved to sustain efficient cardiac performance at rest, during exercise and, importantly, to respond to pathologic stressors ^17^. As the Ca^2+^ responsive element, the CTF is the central sensor and modulator of myofilament activation. It is also a prominent arbiter of molecular relaxation via the release of Ca^2+^ from Site II of cTnC, and the PKA-mediated phosphorylation of Ser 23/24 (S_23_S_24_) within the N-terminus of the inhibitory subunit, cTnI ^18,19^. The phosphorylation of cTnI is known to increase the rate of Ca^2+^ dissociation (*k_off_*) from cTnC and represents a nodal link whereby the myocellular β-adrenergic signaling cascade regulates the ability of the heart to efficiently relax in response to hemodynamic challenges on a beat-to-beat basis ^20^. While the modulation of this dynamic process is dependent on an array of protein-protein interactions within the highly allosteric CTF, the primary biophysical mechanism at the level of the myofilament remains poorly understood, largely due to the lack of extant structure in the N-terminus of cTnI ^21,22^.

In the current study, we utilize an integrated *in vivo – in vitro – in silico* approach to delineate the structural and dynamic basis for mutation-specific alterations in the myofilament response to the β-adrenergic mediated initiation of relaxation at the level of the cardiac sarcomere. We assessed this proposed mechanism for differential early-onset diastolic dysfunction in two independent cTnT-linked HCM mouse models. TR-FRET of fully reconstituted CTFs coupled to molecular dynamics simulations (MD) revealed novel mutation-specific structural refinements of the crucial interface between the N-terminal cTnI phosphorylation loop (cTnI-S_23_S_24_) and Site II of cTnC. These allosterically mediated structural changes are predicted to decrease the access of PKA to cTnI-S_23_S_24_, leading to a decrease in phosphorylation potential, a decrease in the rate of Ca^2+^ dissociation from Site II and a blunted β-adrenergic response *in vivo*. We propose that this complex molecular mechanism underlies the development of early-onset diastolic dysfunction in a subset of HCM patients and may identify a specific target for therapeutic modulation.

## METHODS

### Data Availability

All supporting data, expanded methods, and the major resources table can be found in the Supplementary Material.

## RESULTS

### R92L-cTnT exhibits early-onset diastolic dysfunction prior to hypertrophic remodeling

Impaired relaxation and subsequent diastolic dysfunction have been shown to be among the earliest disease manifestations in sarcomere positive HCM often preceding LV hypertrophy and clinical symptoms. We assessed the time-course of diastolic dysfunction and cardiac remodeling at early (1-2 months) and late (4-6 months) time points in two well-characterized cTnT-linked models of HCM via 2D-echocardiography (Figure 1). While both R92L-cTnT and Δ160E-cTnT are associated with hyper-systolic function from an early age, only R92L-cTnT exhibits an early, progressive decrease in E/e’ driven by a persistent decrease in mitral annulus velocity (Figure 1A-E). In addition, the early, impaired relaxation for R92L-cTnT is not accompanied by an increase in LV wall thickness (T_d_) suggesting that this change is not associated with downstream myocellular responses (Figure 1F). Indeed, once hypertrophy manifests between 4-6 months, no additive change in E/e’ was observed further indicating that hypertrophy is not driving the early-onset diastolic dysfunction observed for R92L-cTnT. Thus, we suspect that for R92L-cTnT but not Δ160E-cTnT there is a proximal, myofilament component that may explain the earlier presentation of diastolic dysfunction.

**Figure 1.**
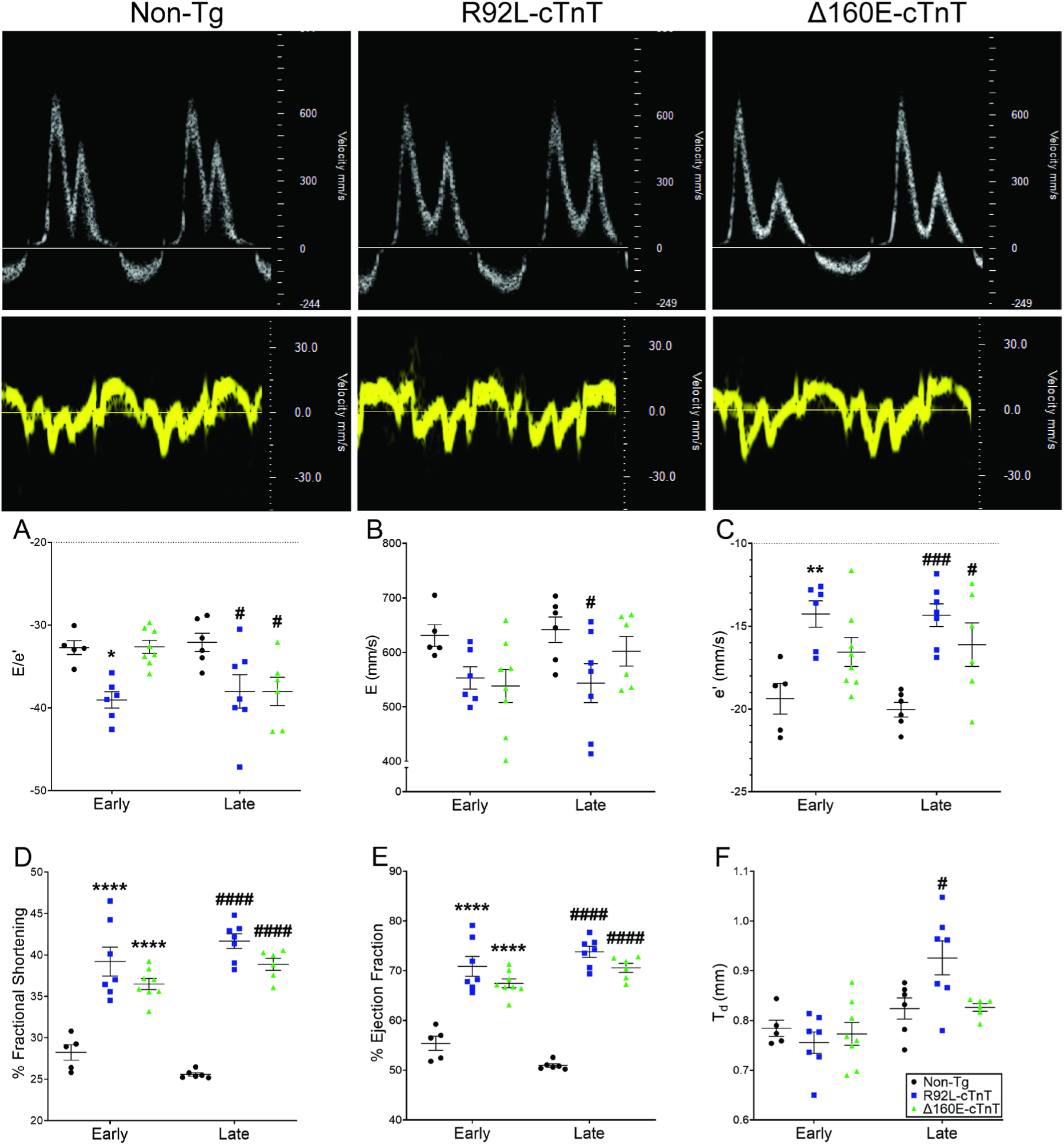
Cardiac function assessed via 2D echocardiography at early (1-2 months) and late (4-6 month) time points in Non-Tg and cTnT-linked models of HCM (R92L-cTnT, Δ160E-cTnT). Top panels show representative traces of the pulsed-wave and tissue doppler at 4-6 months of age. A: Diastolic function assessed by E/e’. B: Early diastolic transmittal flow velocity (E, mm/s). C: Early diastolic mitral annulus velocity (e’, mm/s). D: Systolic function assessed by % Fractional Shortening and E: % Ejection Fraction. F: Total wall thickness (Td, mm). An n = 5-8 was used, mean ± S.E.M is shown. A 2-way ANOVA with Tukey correction was used as described in the Supplemental Materials. Means and adjusted p-values are presented in Table S1 and S2. * p < 0.05 vs NT Early, ** p < 0.01 vs NT Early, **** p < 0.0001 vs NT Early, # p < 0.05 vs NT Late, ### p < 0.005 vs NT Late, #### p <0.0001 vs NT Late.

### cTnI-S_23_S_24_ does not respond to β-adrenergic stimulus in R92L-cTnT mice

Following activation of β-adrenergic receptors, PKA-mediated phosphorylation events modulate the myofilament and sarcoplasmic reticulum axes to coordinate myocellular contraction and relaxation. cTnI (proximal to the mutations) and PLB (distal to the mutations) are critical regulators of myocellular relaxation via phosphorylation sites at serine-16 on PLB and at residues serine-23 and serine-24 on cTnI (cTnI-S_23_S_24_). Previous quantification of these proteins at baseline demonstrated a decrease in the phosphorylation potential of cTnI, but not PLB in R92L-cTnT mutant mice ^23^. To investigate whether changes in cTnI phosphorylation potential could represent a mutation-specific trigger for the observed early-onset diastolic dysfunction we assessed their phosphorylation status at baseline and following *in vivo* stimulation with isoproterenol (Figure 2A). While no differences were observed in baseline expression levels of cTnI and PLB, both R92L-cTnT and Δ160E-cTnT exhibited a potential for reduced expression in the treated groups, likely attributable to the changes in mass associated with these mutations (Figure S1A and C and Figure S2 A-B). Baseline levels of cTnI phosphorylation are known to be high in mice, here we observed a modest, but not significant, increase in pTnI levels upon isoproterenol stimulation for Non-Tg mice ^24^. Unlike Non-Tg mice, both R92L-cTnT and Δ160E-cTnT mice had reduced pTnI expression and a decreased pTnI/TnI ratio (Figure 2B and Figure S1B). Upon stimulus with isoproterenol, R92L-cTnT did not respond exhibiting a decrease in the pTnI/TnI ratio compared to Non-Tg (Figure 2B). Conversely, while pPLB levels and the pPLB/PLB ratio were similar at baseline for all genotypes, R92L-cTnT maintained responsiveness to isoproterenol increasing pPLB/PLB and pPLB expression (Figure 2C, Figure S1D). This suggests that the blunted β-adrenergic response is not caused by diminished or dysregulated PKA function. To determine whether these observations extended to a model of late-onset diastolic dysfunction, we assessed Δ160E-cTnT transgenic mice. While R92L-cTnT exhibited no cTnI phosphorylation response to isoproterenol, Δ160E-cTnT responded as expected with an increase in pTnI/TnI to Non-Tg levels (Figure 2B). Notably, pPLB/PLB levels for Δ160E-cTnT were decreased after isoproterenol treatment illustrating that other myocellular mechanism(s) are present that could account for the late diastolic dysfunction observed (Figure 2C). Given that cTnI expression levels were normal and PKA function intact, these data suggest that the diminished PKA phosphorylation potential of cTnI in the R92L-cTnT mice could account for the differential onset of diastolic dysfunction.

**Figure 2.**
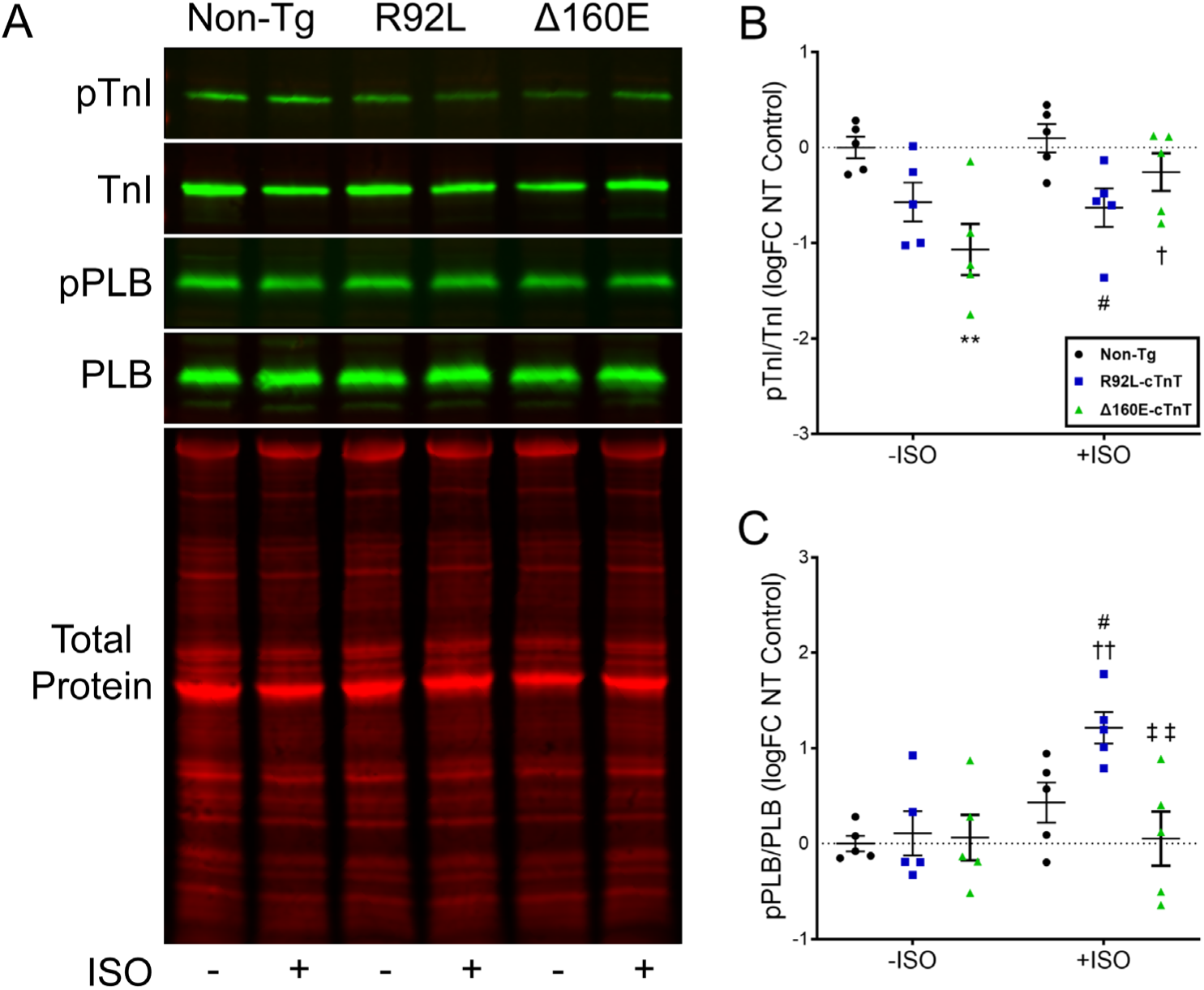
Western blots of phosphorylated cTnI-S_23_S_24_ (pTnI) ratio to cTnI, and the Ser-16 phosphorylation of PLB (pPLB) to total PLB, performed on ventricular homogenates before (-ISO) and following (+ISO) isoproterenol stimulation in Non-Tg, R92L-cTnT, and Δ160E-cTnT mice. A: Representative blots including a total protein stain corresponding to the pTnI blot. B: Summary of pTnI/TnI and C: pPLB/PLB ratios before and following isoproterenol stimulation. Values are reported as mean ± S.E.M., n = 5 animals per group. A 2-way ANOVA with Tukey or Sidak correction was used as described in the Supplemental Materials. Adjusted p-values are presented in Table S3. ** p < 0.01 vs Non-Tg within -ISO; # p <0.05, ## p < 0.01 vs Non-Tg within +ISO; † p < 0.05, †† p < 0.01 vs -ISO within genotype; ‡‡ p < 0.01 vs R92L-cTnT within +ISO.

### Constitutive phosphorylation of cTnI-S_23_S_24_ increases the rate of relaxation in R92L-cTnT hearts

To assess the potential for functional recovery of the early diastolic dysfunction observed in R92L-cTnT hearts via restoration of phosphorylation at cTnI-S_23_S_24_, we employed an *in vivo* phosphomimetic approach utilizing transgenic mice in which the native serines have been replaced with aspartic acids (cTnI-D_23_D_24_) ^25^. Non-transgenic (Non-Tg), R92L-cTnT, and Δ160E-cTnT mice were bred with cTnI-D_23_D_24_ mice to produce Non-Tg/cTnI-D_23_D_24_ (NTDD), R92L-cTnT/cTnI-D_23_D_24_ (RLDD), and Δ160E-cTnT/cTnI-D_23_D_24_ (ΔDD) transgenic mice. We measured cardiac inotropic (+dP/dt) and lusitropic (-dP/dt) responses at baseline and following increasing bolus doses of dobutamine using the Langendorff preparation in the isovolumic mode (Figure 3, Table 1). LV contractile function (+dP/dt) was impaired in R92L-cTnT hearts at the two highest dobutamine doses compared to Non-Tg and NTDD (Figure 3A). NTDD also had greater contractile function at the highest dobutamine dose as compared to Non-Tg and RLDD. RLDD hearts exhibited a similar response as Non-Tg at all doses (Figure 3A). LV relaxation (- dP/dt) in R92L-cTnT mice was impaired at every dobutamine dose when compared to Non-Tg, NTDD, and RLDD (Figure 3B). Of note, the observed diastolic function of RLDD hearts was largely indistinguishable from either Non-Tg or NTDD. To assess whether NTDD was acting as a simple gain of function variant, we repeated these studies on Δ160E-cTnT mice (Figure 3C-D). Similar to R92L-cTnT, Δ160E-cTnT demonstrated impaired contractile function at the two highest doses of dobutamine compared to Non-Tg and NTDD (Figure 3C). In contrast to RLDD, ΔDD hearts showed no improvement. LV relaxation in Δ160E-cTnT was again impaired at every dose compared to Non-Tg and NTDD (Figure 3D). Unlike RLDD, ΔDD only exhibited improvement below 0.1 µmol/L dobutamine when compared to Δ160E-cTnT alone. In addition, ΔDD demonstrated impaired relaxation at levels similar to Δ160E-cTnT at the highest doses of dobutamine. Thus, the cTnI-D_23_D_24_ phosphomimetic ameliorated the impaired relaxation observed in R92L-cTnT but had no significant effect on diastolic performance in Δ160E-cTnT mice. Taken together these observations support a mutation-specific mechanism with proximal myofilament involvement comprising a critical component of early diastolic dysfunction.

**Figure 3:**
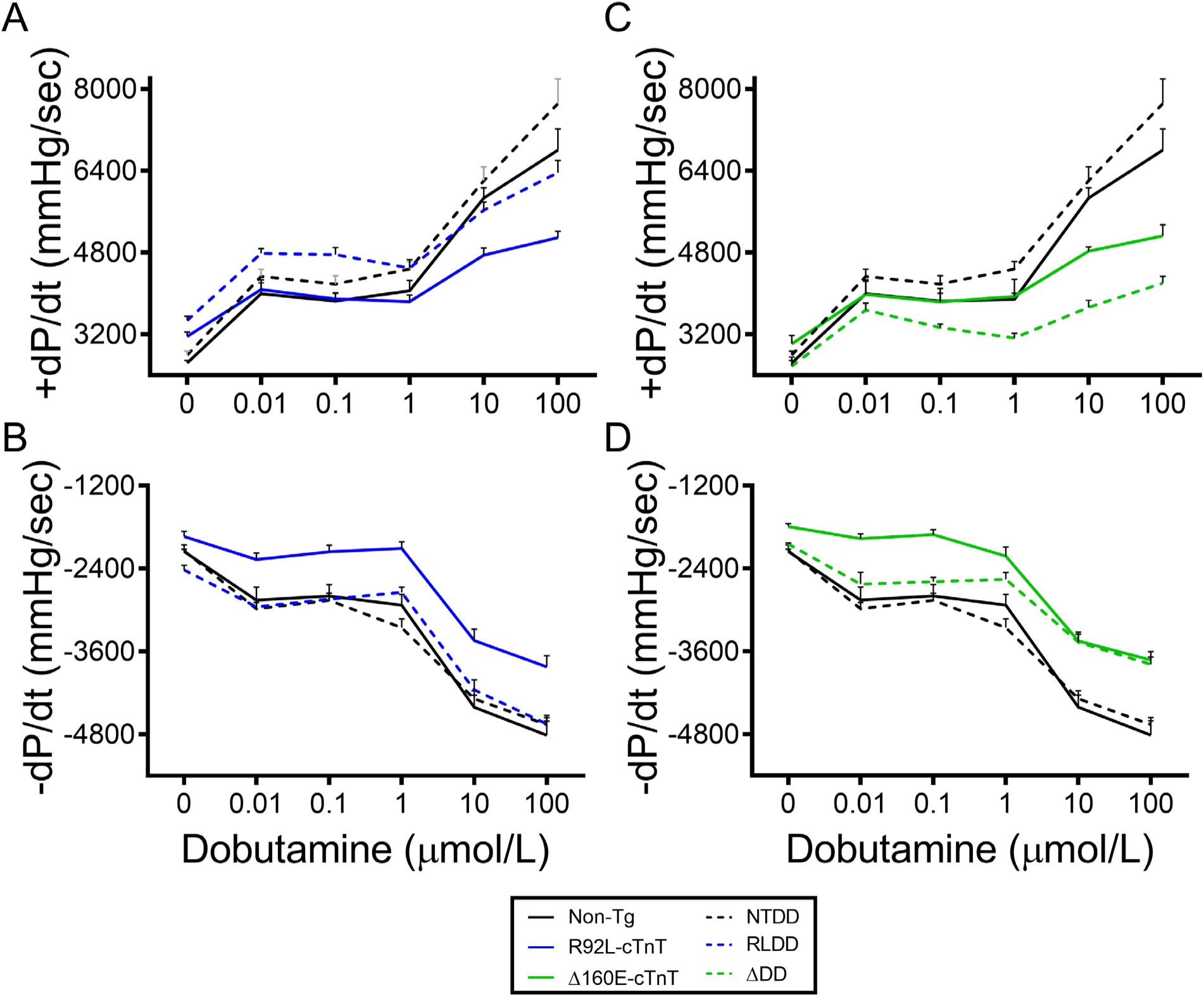
Cardiac inotropic and lusitropic response to dobutamine in isovolumic hearts. A: Maximum +dP/dt and B: minimum –dP/dt in Non-Tg, NTDD, R92L-cTnT, and RLDD. C: Maximum +dP/dt and D: maximum –dP/dt in Non-Tg, NTDD, Δ160E-cTnT, and ΔDD following increasing bolus doses of dobutamine. Values are expressed as means ± S.E.M, n = 4-15 per group. Means and adjusted p-values are presented in Table 1 and Table S4.

**Table 1:** Summary of cardiac inotropic and lusitropic response to dobutamine from Figure 3. Values are reported mean ± S.E.M, n = 4-15 per group. A ROUT method (Q = 10%) was used to remove outliers. A mixed-effects RM 2-way ANOVA with a Tukey or Dunnett correction was used as described in the Supplemental Material. For simplicity a single statistical symbol is shown, adjusted p-values are presented in Table S4. ‖ not significant vs any group, * vs baseline dobutamine dose (0 µmol/L) within genotype, # vs Non-Tg within dobutamine dose, † vs NTDD within dobutamine dose, ‡ vs R92L-cTnT within dobutamine dose, § vs Δ160E-cTnT within dobutamine dose.

| | Non-Tg | NTDD | R92L-cTnT | RLDD | Δ160E-cTnT | ΔDD |
| --- | --- | --- | --- | --- | --- | --- |
| Dobutamine (μmol/L) | Maximum +dP/dt (mm Hg/sec) | | | | | |
| 0 | 2641 ± 47 | 2796 ± 76 \|\| | 3156 ± 90 #† | 3472 ± 83 #† | 3011 ± 163 \|\| | 2577 ± 177 \|\| |
| 0.01 | 3994 ± 270 * | 4330 ± 139 * | 4077 ± 127 * | 4782 ± 90 *‡ | 3977 ± 261 * | 3674 ± 135 *† |
| 0.1 | 3846 ± 154 * | 4183 ± 164 * | 3890 ± 118 * | 4755 ± 142 *#‡ | 3834 ± 259 * | 3329 ± 69 † |
| 1 | 4051 ± 204 * | 4475 ± 147 * | 3840 ± 129 *† | 4494 ± 168 *‡ | 3942 ± 333 * | 3125 ± 96 #† |
| 10 | 5865 ± 196 * | 6204 ± 268 * | 4747 ± 141 *#† | 5628 ± 155 *‡ | 4822 ± 85 *#† | 3727 ± 135 #†§ |
| 100 | 6800 ± 415 * | 7712 ± 480 * | 5090 ± 127 *#† | 6362 ± 235 *‡ | 5391 ± 329 *#† | 4201 ± 132 *#†§ |
| Dobutamine (μmol/L) | Maximum -dP/dt (mm Hg/sec) | | | | | |
| 0 | -2154 ± 93 | -2145 ± 22 \|\| | -1937 ± 72 \|\| | -2418 ± 65 †‡ | -1793 ± 43 #† | -2047 ± 16 §† |
| 0.01 | -2858 ± 191 * | -2983 ± 92 * | -2270 ± 96 *† | -2956 ± 99 *‡ | -1970 ± 73 #† | -2628 ± 173 * |
| 0.1 | -2798 ± 166 * | -2861 ± 101 * | -2159 ± 95 *#† | -2843 ± 87 *‡ | -1909 ± 72 #† | -2590 ± 61 § |
| 1 | -2931 ± 158 * | -3258 ± 128 * | -2108 ± 93 *#† | -2745 ± 75 *†‡ | -2219 ± 131 *#† | -2557 ± 97 † |
| 10 | -4407 ± 174 * | -4284 ± 113 * | -3442 ± 166 *#† | -4155 ± 143 *‡ | -3447 ± 123 *#† | -3466 ± 115 *#† |
| 100 | -4815 ± 208 * | -4660 ± 101 * | -3824 ± 161 *#† | -4649 ± 123 *‡ | -3724 ± 123 *#† | -3789 ± 106 *#† |

### cTnI-D_23_D_24_ increases the rate of Ca^2+^ dissociation for R92L-cTnT filaments

Previous studies have shown that phosphorylation of cTnI-S_23_S_24_ increases the rate of cardiac muscle relaxation^26^. Specifically, stopped-flow Ca^2+^ dissociation kinetics revealed that incorporation of the phosphomimetic cTnI-D_23_D_24_ increases the rate of CTF deactivation identifying this property as a primary molecular mechanism for increased lusitropy ^20^. To begin to assess a potential myofilament-based mechanism to address our *in vivo* results, we utilized stopped-flow Ca^2+^ dissociation kinetics in fully reconstituted CTFs based on the human sequences (Figure 4) ^27^. Wild type (WT) cTnT, R92L-cTnT, and Δ160E-cTnT were used along with WT-cTnI or cTnI-D_23_D_24_ as described in the Supplemental Materials producing WT-cTnT/cTnI-D_23_D_24_ (WTDD), R92L-cTnT/cTnI-D_23_D_24_ (RLDD), and Δ160E-cTnT/cTnI-D_23_D_24_ (ΔDD) CTFs. Both R92L-cTnT and Δ160E-cTnT induced a decrease in the rate of Ca^2+^ dissociation compared to WT consistent with previous studies of HCM-linked mutations (WT 117.0 ± 0.66 s^-1^, R92L-cTnT 83.47 ± 1.00 s^-1^, Δ160E-cTnT 82.17 ± 1.553 s^-1^) (Figure 4B) ^28^. To determine whether the incorporation of the phosphomimetic cTnI-D_23_D_24_ ameliorates impaired CTF deactivation in a mutation-specific manner, we assessed the rates for WTDD, RLDD and ΔDD. We should note that it was necessary to run the ΔDD kinetics at low ionic strength in order to fit an exponential decay function likely due to the known primary effect of the Δ160E-cTnT on weak actin-myosin binding (Figure S3) ^29^. The addition of cTnI-D_23_D_24_ to the WT CTF induced the expected increase in the rate of Ca^2+^ dissociation (WTDD 587.17 ± 18.10 s^-1^) (Figure 4B). The reduction in the rate of Ca^2+^ dissociation induced by R92L-cTnT was absent in RLDD filaments which exhibited a rate similar to WTDD (RLDD 529.4 ± 1.21 s^-1^). To assess whether this result was specific to R92L-cTnT, we measured dissociation rates in ΔDD. Unlike RLDD, the phosphomimetic did not induce a change sufficient to return to WTDD rates, with ΔDD having a decreased Ca^2+^ dissociation rate compared to WTDD (ΔDD 496.8 ± 23.68 s^-1^) (Figure 4B). Taken together, the early-onset diastolic dysfunction and mutation-specific decrease in phosphorylation potential associated with R92L-cTnT coupled to the observed functional improvements upon genetic incorporation of cTnI-D_23_D_24_ supports a potential interaction between the N-lobe of cTnC and the N-terminus of cTnI that can be allosterically modulated by cTnT.

**Figure 4:**
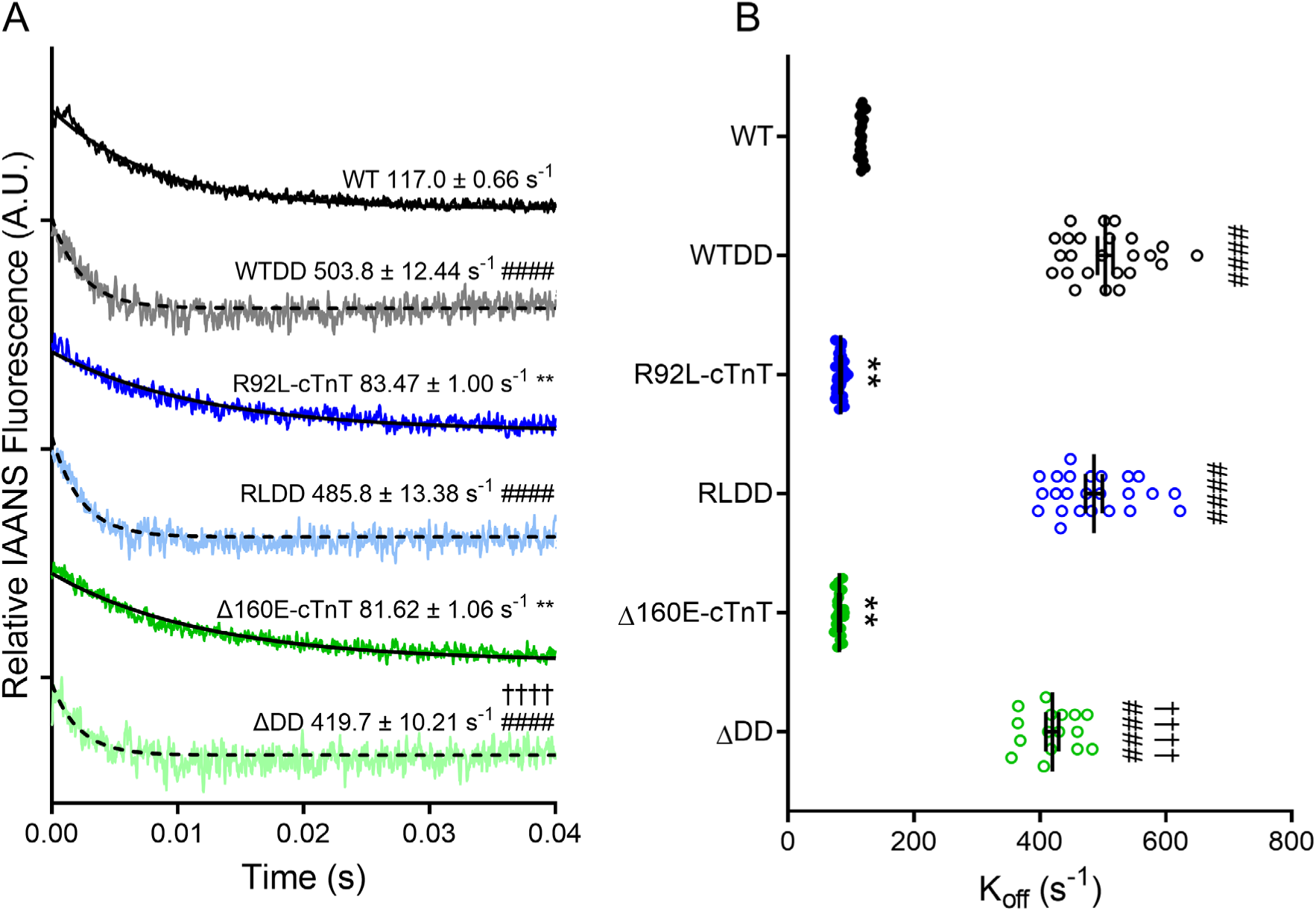
Ca^2+^ dissociation rates from reconstituted CTFs. A: Representative traces. B: Rate of Ca^2+^ dissociation in WT, WTDD, R92L-cTnT, RLDD, Δ160E-cTnT, and ΔDD CTFs. Values are expressed as mean ± S.E.M. A 2-way ANOVA with Sidak correction for MC was used as described in the Supplemental Materials. ** p < 0.01 vs WT, #### p < 0.0001 vs baseline (without phosphomimetic) within genotype, †††† p < 0.0001 vs WTDD. Adjusted p-values are presented in Table S5.

### Determining the structure of the WT cTnC-cTnI interface

In order to better understand the molecular basis for the observed functional coupling of the cTnI-S_23_S_24_ phosphorylation state and cTnC Ca^2+^ dissociation kinetics at Site II, we developed a TR-FRET based approach utilizing a fully reconstituted CTF (based on the human sequences) to directly determine the structural interactions between these two domains. Despite recent advancements in cryo-electron microscopy and studies that have resolved the bulk structure of the CTF, the N-terminus of cTnI (containing S_23_S_24_) has yet to be resolved due to its inherent flexibility ^21,22^. To determine this central structure, we measured the distances (Å) and full-width half-max (FWHM, Å) of the distance distributions (a measure of variability of these distances). Taken together, these measurements provide the range of possible changes in distance associated with environmental or structural alterations to the CTF. Three TR-FRET pairs were chosen to span the cTnI N-terminus: cTnC84C-cTnIA9C, cTnC84C-cTnIA17C, and cTnC84C-cTnIA28C, or A9C, A17C and A28C, respectively (Figure 5). At site A9C we measured a distance of 52.9±2.114 Å with a FWHM of 22.3±1.157 Å with no change upon the addition of saturating Ca^2+^ (48.3±1.143 Å, 21.2±1.026 Å) (Figure 5A, Table S6). The more distal amino acids at sites A17C and A28C exhibited a decrease in distance at baseline (43.2 ±0.868 Å, 41.8± 0.620Å), indicating that these regions of the N-terminus of cTnI are closer to cTnC (Figure 5B and C, Table S4). This decrease in distance was accompanied by a decrease in FWHM (17.8± 1.113 Å to 14.8 ±1.249 Å), consistent with a more ordered state and greater interactions with cTnC. Additionally, site A17C showed a decrease in distance (43.2 ±0.868 Å to 39.1±0.534Å) upon the addition of saturating Ca^2+^. These results support the conclusion that the presence of Ca^2+^ results in more interactions between the N-terminus of cTnI and cTnC within these domains, establishing a baseline structure for this multiprotein complex of the CTF core in both states.

**Figure 5:**
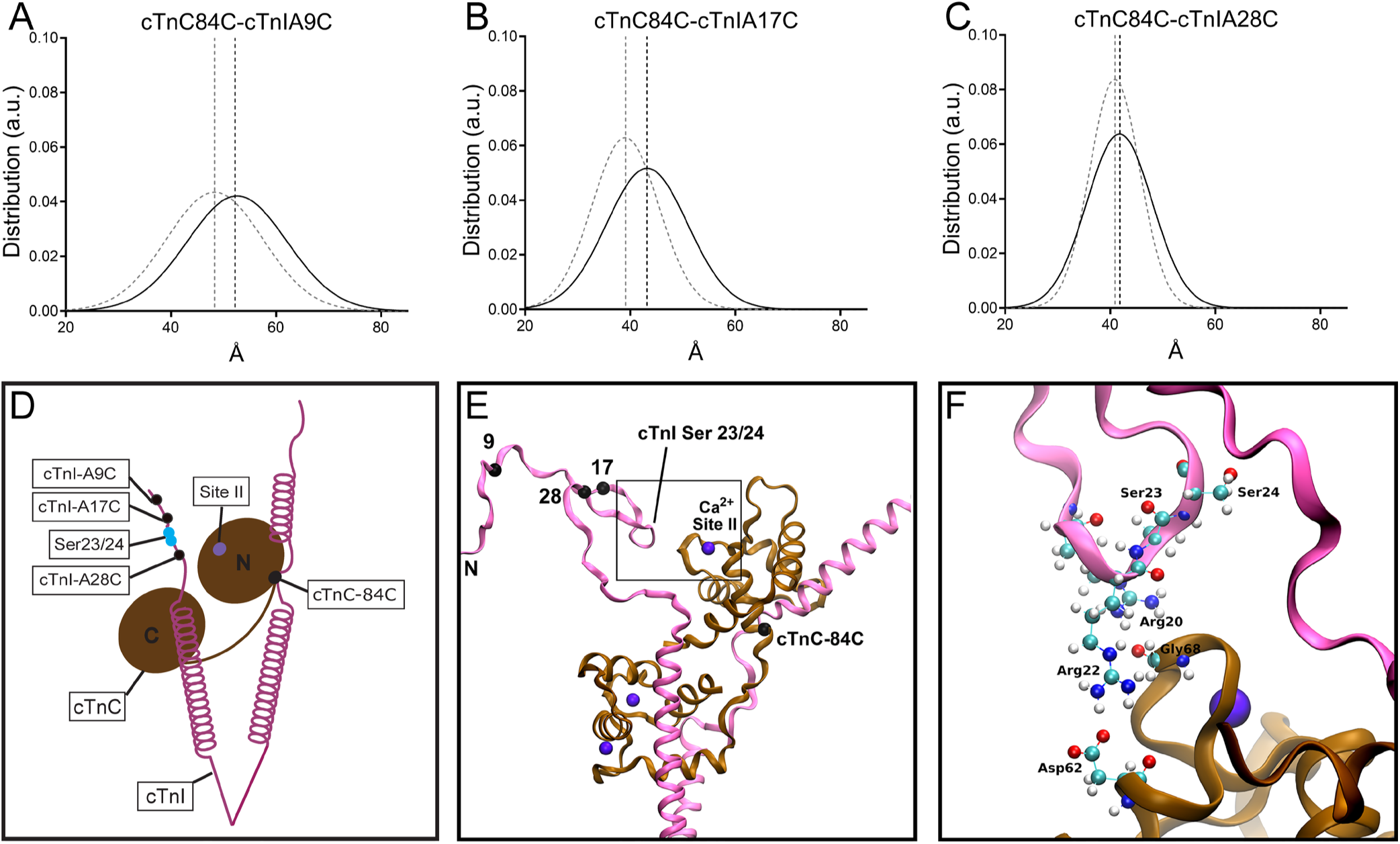
Structure of the WT cTnC-cTnI Interface determined by TR-FRET and MD. Distance distributions of the WT cTnC-cTnI interface generated via TR-FRET between A: cTnC84C to cTnIA9C, B: cTnC84C to cTnIA17C, C: cTnC84C to cTnIA28C. D: Schematic diagram of the cTnC-cTnI interface; cTnC in brown, cTnI in magenta, cTnI-S_23_S_24_ in cyan dots, cTnC Site II in purple dot, and TR-FRET sites in black dots. E: MD of the WT cTnC-cTnI interface labeled as the schematic. F: Zoom-in of box outlined in panel E. Relevant side chains are labeled with their respective side chain amino acid and site (eg. Ser23). Means of the distances and FWHMs obtained and adjusted p-values are presented in Table S6 and S7.

### Atomistic level resolution of the WT cTnC-cTnI interface

To obtain deeper mechanistic insight into the dynamic regulation of the WT cTnC-cTnI interface, we next employed our all-atom model of the CTF (all human sequences) ^30^. A full atomistic structure was first developed by restraining the existing model to the TR-FRET distances identified above followed by the full relaxation of all other atomic positions. This approach generates a structure in which all atoms can adapt to the average distances derived from the experimental values. We then utilized this model to focus on cTnC-cTnI interactions near Site II of cTnC, where the physiological Ca^2+^ binds. The resultant cTnC-cTnI interactions in the computed average structures from the MD are shown in Figure 5D-E. While residues 9, 17, and 28 of cTnI were subject to distance constraints, the remainder of the N-terminus of cTnI was free to search for electrostatic contacts. In the WT system, cTnI anchors slightly below cTnC site II with two arginine residues, 20 and 22. Arg20-cTnI comes within electrostatic contact (< 5 Å) of the backbone carbonyl group of Gly68-cTnC. Arg22-cTnI reaches further underneath cTnC site II, creating a salt bridge with Asp62-cTnC. The breadth of the FWHM of the distance distributions of the TR-FRET signals in the +Ca^2+^ condition at A17C suggests that these interactions are transient. Thus, with respect to the nearby PKA-mediated phosphorylation sites, in the WT state, cTnI-S_23_S_24_ is clearly solvent exposed and accessible for PKA-mediated phosphorylation in the context of β-adrenergic activation.

### Structural effects of R92L-cTnT and Δ160E-cTnT on the cTnC-cTnI interface

To directly test the hypothesis that the presence of R92L-cTnT results in a decrease in physical access to the PKA consensus sequence, we again utilized TR-FRET. Two TR-FRET pairs were chosen to flank the consensus sequence of PKA, cTnC84C-cTnIA17C and cTnC84C-cTnIA28C, A17C and A28C respectively. At site A17C, the addition of Ca^2+^ reduces the cTnC-cTnI interface distance by approximately 4Å for WT CTFs (43.2±0.868 Å to 39.1±0.534 Å), with no concomitant change in FWHM (18.2±1.303 Å to 14.9±0.669 Å), (Figure 6A-C, Table S8). Similarly, at site A17C, both the R92L-cTnT and Δ160E-cTnT displayed a decrease in distances upon the addition of Ca^2+^ of similar magnitude (Figure 6A, Table S8). However, only for R92L-cTnT was a decrease in FWHM observed at A17C compared to WT (11.7±1.709 Å) indicating that when Ca^2+^ activates R92L-cTnT filaments the distribution of available distances is reduced, thus constraining the cTnC-cTnI interface in its closest conformation. Unlike A17C, at site A28C, the data reveals that the addition of Ca^2+^ does not affect the distance of the cTnC-cTnI interface for WT filaments (41.8±0.620 Å and 40.9±0.428 Å) (Figure 6D-F, Table S8). For R92L-cTnT, however, with the addition of Ca^2+^, we observed a decrease in distance of approximately 3Å compared to R92L-cTnT at baseline (from 42.5±0.715 Å to 39.7±0.680 Å) with a concomitant decrease in FWHM compared to WT at baseline (11.1±0.965Å) (Figure 6D-E). These changes were not observed with Δ160E-cTnT filaments. Consistent with our observations at site A17C, these data indicate that upon Ca^2+^ activation, site A28C is closer to Site II of TnC, in proximity to the regions directly flanking the PKA recognition sequence. Of note, a small but not significant drop in FWHM at site A28C occurs for WT filaments (from 14.7±1.177 Å to 11.2±0.601 Å) while Δ160E-cTnT displays the opposite, sustaining the FWHM upon the addition of Ca^2+^ (from 15.4±1.219 Å to 14.6±0.899 Å). Taken together, these observations suggest an allosterically mediated structural mechanism whereby the R92L-cTnT mutation alone disrupts the accessibility of PKA to its primary cTnI recognition sequence, resulting in a decrease in cTnI-S_23_S_24_ phosphorylation potential as we observed *in vivo* (Figure 2B).

**Figure 6:**
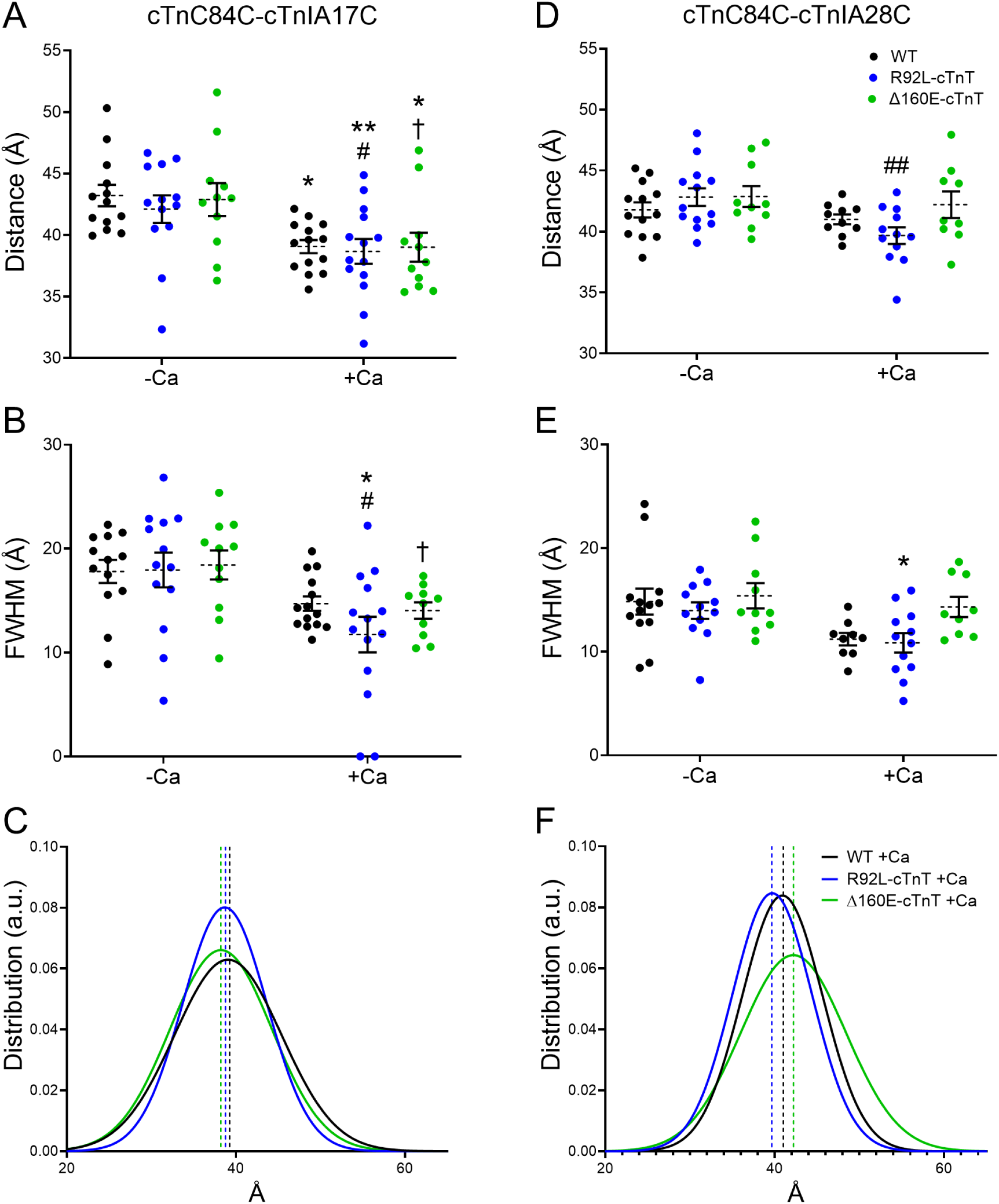
TR-FRET flanking the PKA recognition sites in CTFs for WT, R92L-cTnT, and Δ160E-cTnT. For TR-FRET pair cTnC84C-cTnIA17C - A: Distances (Å), B: FWHM (Å), and C: Averages distance distribution plots. For FRET pair cTnC84C-cTnIA28C - D: Distances (Å), E: FWHM (Å), and F: Averages distance distribution plots. Values are expressed as mean ± S.E.M., n = 9-14 per group. A 2-way ANOVA with Tukey correction was used as described in the Supplemental Materials. Means of distances and FWHMs obtained and adjusted p-values are presented in Table S8 and S9. * p< 0.05, ** p<0.01 vs WT -Ca; # p<0.05, ## p<0.01 vs R92L-cTnT -Ca; †p<0.05 vs Δ160E-cTnT -Ca.

### Atomistic level resolution of the R92L-cTnT and Δ160E-cTnT cTnC-cTnI interface

MD incorporating the R92L-cTnT mutation demonstrated a repositioning of the cTnC-cTnI interactions around cTnC Site II (Figure 7A-B). The N-terminus of cTnI lies close to cTnC Site II, shifting slightly closer as compared to the WT structure. While Arg22-cTnI retains the interaction with Asp62-cTnC, it gains a novel contact to the backbone carbonyl of cTnC62. Arg22-cTnI also shifts farther underneath Site II, interacting with the backbone carbonyl of cTnC65. These interactions are due to Arg20-cTnI shifting away from cTnC and orienting towards the solvent, losing the contact previously present in the WT structure, and creating space for Arg22-cTnI to move closer to cTnC. Additionally, the R92L-cTnT mutation brings cTnI-S_23_ into contact with cTnC site II, residing between the backbone carbonyls of residues Glu66 and Asp67 of cTnC. cTnI-S_24_ remains solvent exposed. Together these data support that the structure of the consensus sequence is changed in R92L-cTnT myofilaments leading to an allosterically mediated alteration in cTnI phosphorylation potential.

**Figure 7:**
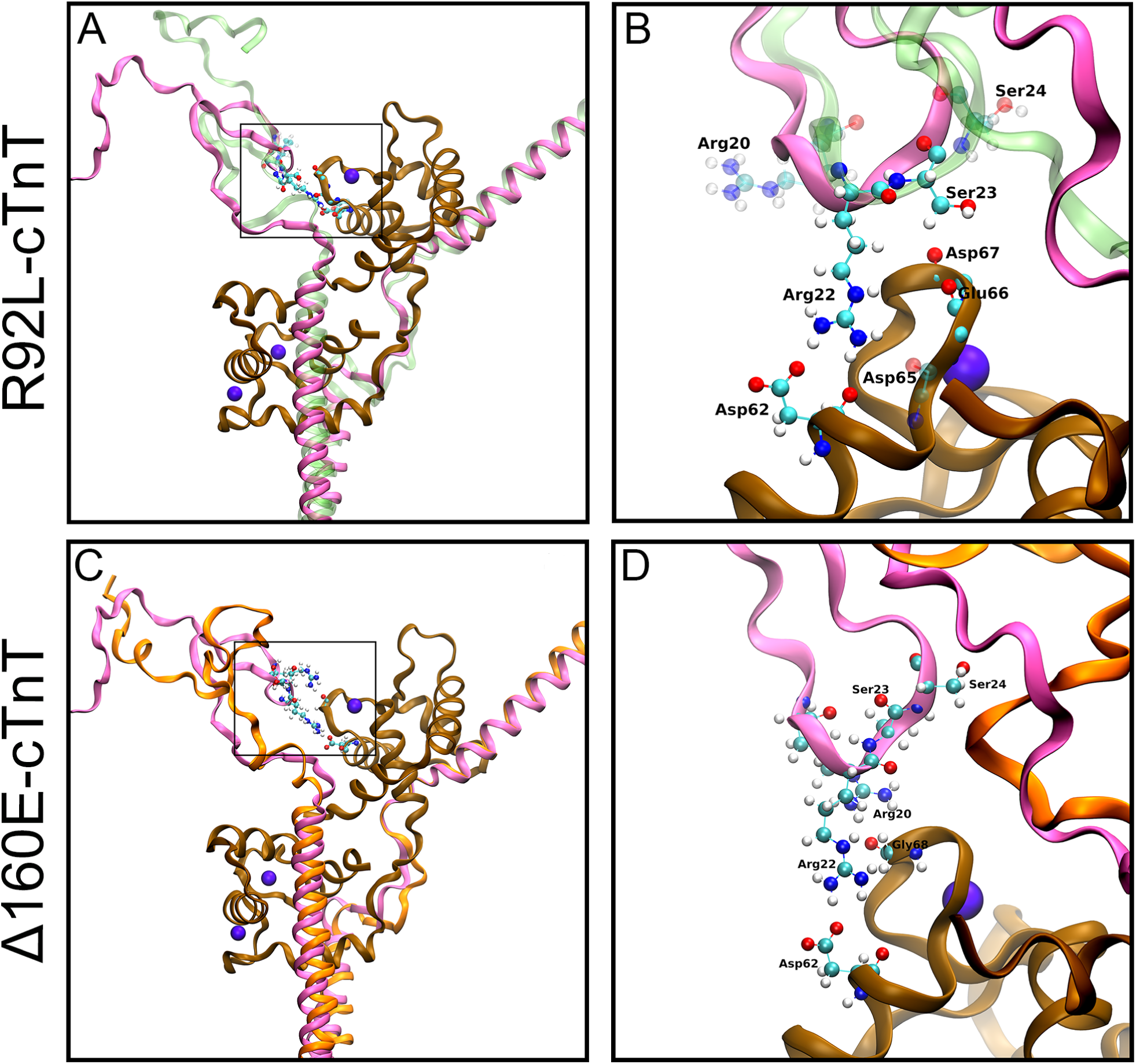
Interactions at the cTnC-cTnI interface for R92L-cTnT and Δ160E-cTnT produced via MD. A: MD of the R92L-cTnT cTnC-cTnI interface as in Figure 4B with the R92L-cTnT overlayed in green over the WT (magenta). B: Zoom-in of the box outlined in panel A. Side chain interactions shown are those altered by the inclusion of R92L-cTnT. C: MD of the Δ160E-cTnT cTnC-cTnI interface with the Δ160E-cTnT overlayed in orange. D: Zoom-in of the box outlined in panel C. Side chain interactions shown are those of WT as no interactions are present for Δ160E-cTnT.

Conversely, the Δ160E-cTnT structure reveals that cTnI shifts towards the solvent, away from cTnC (Figure 7C-D). This change in the cTnC-cTnI interface structure is substantial as all previous cTnC-cTnI interactions present in the WT structure are completely absent (Figure 7D). cTnC-cTnI interactions are likely still present due to the inherent flexibility of the N-terminus of cTnI as demonstrated by the FWHM of the FRET distance distributions. However, they are likely more transient than the cTnC-cTnI interactions found in the WT and R92L-cTnT structures. The electrostatic interactions observed with R92L-cTnT but not with Δ160E-cTnT are likely the cause of the decreased FWHM observed in TR-FRET for R92L-cTnT. Together this results in a more constrained conformation of the N-terminus of cTnI that likely attenuates PKA’s access to its consensus sequence and blunts cTnI phosphorylation.

## DISCUSSION

While the classic definition of HCM is deceptively simple, the modern clinical reality is far more complex as phenotypic variability is not uncommon. Contributors to phenotypic variability are often secondary responses to the primary sarcomeric mutation, including activation of downstream signaling cascades, environmental factors, co-morbidities, male sex and genetic modifiers to name a few ^31^. Given that HCM patients are often diagnosed later in life when symptoms develop, and patient tissue samples are usually obtained late-stage in the context of surgical myectomy the primary mechanistic linkage between genotype and phenotype is difficult to ascertain as the earliest stages of disease pathogenesis are simply not accessible. An array of murine models have been developed to provide insight into this crucial pathogenic stage and longitudinal progression in an intact system ^32^. Here we utilized two independent and well-characterized transgenic models of cTnT-linked HCM based on known human mutations, R92L-cTnT (missense) and Δ160E-cTnT (single amino acid in-frame deletion). Both represent well-established mutational hotspots, reside in distinct functional domains, disrupt CTF structure/dynamics, and exhibit different phenotypes in patients and our mice. While both R92L-cTnT and Δ160E-cTnT mice exhibit diastolic dysfunction and a similar degree of Ca^2+^ hypersensitivity of force development in skinned fibers, secondary patterns of cardiac remodeling *in vivo* are markedly distinct with Δ160E-cTnT alone exhibiting extensive fibrosis and disruption of downstream myocellular Ca^2+^ homeostasis ^23,33,34^. This clear difference in disease progression suggests a differential mechanism likely exists to explain this variability. Indeed, despite their similarities, manifestations of early impaired relaxation observed as a marked reduction in E/e’, driven primarily by a loss of mitral annulus velocity, were detected in R92L-cTnT mice in the absence of overt cardiac remodeling (Figure 1). Thus, it is likely that the different time-of-onset of diastolic dysfunction observed represents a mutation-specific primary myofilament mechanism at the level of the myofilament that underlies a subset of HCM.

While the downstream targets of the PKA-mediated activation of cardiac performance are broad, two central components are the regulation of Ca^2+^ homeostasis and flux (largely SR-mediated) and the myofilament (cTnI, MyBP-C and Titin). In the context of heart failure, hypo-phosphorylation of PLB Ser-16 (inhibition of SR Ca^2+^ uptake) and cTnI-S_23_S_24_ (failure to decrease Ca^2+^ sensitivity) are consistent findings and usually correlated with late-stage disease and a blunted response to β-adrenergic activation ^35^. From the standpoint of sarcomeric HCM, a decrease in phosphorylation potential has been observed for both the myofilament and SR-mediated regulatory components ^36,37^. A study by Najafi et al. utilized a unique set of mutant MyBPC-3 mice (in both the homozygous and heterozygous states) carrying a known HCM-linked point mutation to assess whether a component of the pathogenic profile could be linked to a disruption of the proposed balance of phosphorylation between these two pathways ^24^. An important corollary of that study was the observation that the PKA-mediated phosphorylation potential of PLB was preserved despite a decrease in cTnI phosphorylation (similar to our observation in the R92L-cTnT mice). Fraysse et al., using the same models, then suggested that diastolic dysfunction (and myofilament Ca^2+^ sensitization) may represent an earlier phenotypic presentation independent of overt remodeling ^38^. In the current study, we have shown that R92L-cTnT exhibits early diastolic dysfunction in the absence of an increase in wall thickness (Figure 1). Similarly, we detected a non-uniform blunting of PKA-mediated phosphorylation of its targets localized to cTnI for R92L-cTnT (Figure 2B). Conversely, while Δ160E-cTnT maintained its cTnI PKA responsiveness it exhibited a downstream disruption in PLB response to stimulation likely related to the later onset of diastolic dysfunction. Based on these observations, we propose that the diminished PKA phosphorylation potential of cTnI caused by R92L-cTnT accounts for the different time-of-onset of diastolic dysfunction in these two cTnT-linked HCM mouse models.

Alterations in PKA-mediated phosphorylation of cTnI are known to be a critical regulator of diastolic function. Mice with changes to the N-terminus of cTnI that result in an inability to phosphorylate cTnI-S_23_S_24_ (slow skeletal cTnI and alanine substituted S_23_S_24_) exhibit increased Ca^2+^ sensitivity and impairment in cardiac relaxation at baseline and following β-adrenergic stimulation ^20,39-44^. Conversely, phosphomimetic cTnI mice (cTnI-D_23_D_24_) mice demonstrated decreased Ca^2+^ sensitivity, enhanced cardiac contractility and relaxation, and enhanced force-frequency contractility ^45-47^. Based on our observations of early diastolic dysfunction associated with diminished phosphorylation potential of cTnI-S_23_S_24_ we asked if introduction of cTnI-D_23_D_24_ was sufficient to recover function for R92L-cTnT. While R92L-cTnT demonstrated impaired relaxation (-dP/dt) at every dobutamine dose, relaxation was completely restored to Non-Tg levels in RLDD; ΔDD showed no functional improvement (Figure 3). Moreover, unlike R92L-cTnT, Δ160E-cTnT mice have impaired downstream Ca^2+^ handling, demonstrated by decreased SR Ca^2+^ uptake and increased NCX protein levels. The cTnI-D_23_D_24_ substitution cannot fully restore function in Δ160E-cTnT mice to the level observed in R92L-cTnT mice^34,48^. Thus, with PKA function intact, and the restoration of the diastolic response to incorporation of cTnI-D_23_D_24_, we propose that the early diastolic dysfunction associated with R92L-cTnT represents a primary myofilament-based mechanism affecting the complex inherent interplay between PKA-mediated cTnI phosphorylation and Ca^2+^ sensitivity that is the central molecular arbiter of diastolic performance.

Since the initial results of Solaro, et al., demonstrating a kinetic link between PKA-mediated cTnI phosphorylation and a decrease in the Ca^2+^ sensitivity of myofilament activation in the intact heart, the primary mechanisms involved in this pivotal physiologic response have been extensively studied ^49^. In the R92L-cTnT mouse model, previous work has suggested that the impaired β-adrenergic response is isolated to the myofilament axis because downstream Ca^2+^ dynamics were intact ^23,33,50-53^. The development of the TnC-T53C probe has greatly improved our ability to investigate mutation-specific changes to Ca^2+^ binding profiles, allowing us to observe changes to Ca^2+^ dissociation rates rather than steady-state measurements ^54,55^. We observed a reduction in dissociation rates for both R92L-cTnT and Δ160E-cTnT, illustrating that while the downstream Ca^2+^-axis appears to be intact for R92L-cTnT, the mutations alone are sufficient to alter Ca^2+^ kinetics at the myofilament (Figure 4). Notably, Davis et al. illustrated that many HCM-causative mutations result in a decrease in CTF dissociation rates in this isolated *in vitro* environment ^28^. This suggests that independent of the coupling of Ca^2+^ sensitivity to TnI phosphorylation, additional structural mechanisms exist which can alter CTF dissociation rates. These mechanisms may be mediated by changes in the inhibitory region or switch peptide which we did not assay in the current study. Upon incorporation of cTnI-D_23_D_24_ we found that for R92L-cTnT, but not Δ160E-cTnT, there is a recovery of dissociation rates towards WT levels (Figure 4). This suggests that for R92L-cTnT, structural changes at the cTnC-cTnI interface, capable of altering Ca^2+^ dissociation from cTnC and accessibility of the PKA consensus sequence, could represent a mutation-specific nodal point in disease mechanism. In fact, phosphorylation of cTnI-S_23_S_24_ has been shown to weaken interactions between cTnC and cTnI ^56^. These weakened interactions may be responsible for the enhanced increase in Ca^2+^ dissociation observed with phosphomimetic cTnI-D_23_D_24_. We previously proposed that structural changes to the N-terminus of cTnI are responsible for differential Ca^2+^ dissociation rates observed in HCM mutations ^57^. These studies suggest that in both the context of health and disease these interactions are crucial for regulating Ca^2+^ dissociation; however, our significant structural refinement of the cTnC-cTnI interface was required to make definitive conclusions.

Current structural models of the CTF underscore the inherent flexibility of multiple regions within the WT complex, that have long complicated the ability to generate a complete structural model ^21,22,58-60^. Multiple inherently flexible regions of the complex are critical in regulating CTF function including the N- and C-terminus of cTnI, the linker region of cTnT, and the N- and C-terminus of cTnT. Resolving these regions is important for understanding the biophysics that underlie the structure-function relationship of the WT CTF. In the present study, our TR-FRET and MD have further elucidated the structure of the N-terminus of cTnI, placing it in proximity of the N-lobe of cTnC (Figure 5A-C). The structure of the N-terminus of cTnI was initially studied with fragments using NMR. Finley et al. showed that the cTnI fragment 1-80 makes contacts with the N-terminus and C-terminus of cTnC ^61^. The general observation that the N-terminus of cTnI interacts with the N-lobe of cTnC has been recapitulated in multiple systems spanning from fragment studies to the troponin complex alone ^56,62-65^. This study represents the first time the N-terminus of cTnI has been resolved in fully reconstituted CTFs. Of note, our observation that the extreme N-terminus of cTnI (at A9C) is more disordered than the more C-terminal regions (A17C and A28C) independent of Ca^2+^ status is consistent with previous work, validating our approach ^56^. Recently, Pavadai et al used protein docking and MD to further elucidate the N-terminal cTnC-cTnI interface in another CTF model ^60^. Despite the inherent differences in these two models, they both show that Ca^2+^ saturated CTFs show significant interactions between cTnI-R20 and cTnI-R22 and cTnC, in agreement with our MD results (Figure 5D-F). While some of the specific interactions highlighted differ from ours, possibly owing to their exclusion of unstructured regions of the molecule, the observation of interactions between cTnI and cTnC remain consistent.

The ability of point mutations within these flexible regions to affect the structure of distant regions of the complex is an ongoing topic of study. We have previously demonstrated that single amino missense mutations at the 92 codon “hotspot” of the TNT1 domain of cTnT cause distinct alterations to peptide dynamics that result in structural changes ^33,53^. Specifically, MD in R92L-cTnT demonstrated alterations to secondary structure including pronounced hinge motion resulting in distinct oscillations surrounding residue 104 and unfolding of the helical structure across residues 70-170 of cTnT. It is likely that these mutation-specific alterations to secondary structure caused by the R92L-cTnT mutation are propagated within cTnT to the IT arm and disrupt conformational changes within the troponin complex. The IT arm is a coiled-coil in the troponin core where extensive subunit-subunit interactions between cTnI and cTnT make this region the most stable portion of the troponin complex ^66,67^. Given that the rest of the troponin complex is highly flexible, changes to the stability of the IT arm, induced by mutations in cTnT, may alter the allosteric conformational changes that normally occur within the complex. In 2016 Williams et al. demonstrated that R92L-cTnT and R92W-cTnT transmitted structural changes to the IT arm ^57^. This structural alteration could result in a decrease in PKA solvent accessibility to its consensus sequence on cTnI-S_23_S_24_. Collectively, the structural changes we observed in R92L-cTnT via TR-FRET that result in allosterically mediated conformational changes at the cTnC-cTnI interface in regions flanking the PKA consensus sequence, directly confirm these model predictions (Figure 6). Additionally, our MD studies of the cTnC-cTnI interface in the presence of both R92L-cTnT and Δ160E-cTnT support a structural mechanism for the blunted β-adrenergic response that is isolated to cTnI for R92L-cTnT (Figure 7).

In the current study we have utilized an integrated *in vivo – in vitro – in silico* approach to identify a novel allosteric mechanism for early-onset diastolic dysfunction caused by HCM-linked CTF mutations. Decreased LV compliance in patients with HCM is common, multifactorial, and often progressive. It underlies elevated LV filling pressures at baseline and, importantly, contributes to dyspnea on exertion, which significantly impacts quality of life in affected individuals. While multiple secondary myocellular pathways are known to be involved in late-stages of HCM, advanced imaging of pre-clinical cohorts has consistently revealed that impaired relaxation often precedes evidence of structural cardiac remodeling ^12-14^. Thus, to define the primary mechanism of this early-onset diastolic dysfunction, we have focused on the primary molecular regulator of cardiac relaxation, the CTF. Specifically, we compared two independent transgenic mouse models carrying cTnT mutations linked to HCM that exhibit impaired relaxation, diverse patterns of ventricular remodeling and markedly blunted responses to β-adrenergic activation. Our results revealed a distinct, mutation-specific molecular mechanisms including, in one case, an allosteric disruption of the structural and dynamic regulation of the highly coupled PKA-mediated phosphorylation of cTnI and the rate of dissociation of Ca^2+^ from Site II of cTnC. At the atomic level, novel cTnT mutation-induced interactions between cTnI and cTnC are predicted to limit the physical access of PKA to cTnI-S_23_S_24_. These interactions result in a “dominant” decrease in phosphorylation potential and lead to an impairment in the molecular initiation of relaxation. We propose that both the identification of this novel primary mechanism and the tools we have developed to explore the coupling of cTnI phosphorylation to Ca^2+^ dissociation kinetics provide a unique opportunity for screening and eventually modulating this primary pathogenic target.

## LIMITATIONS

We acknowledge the lack of human-tissue derived experimental systems in this work. With the long-term goal of facilitating translational insight both the CTF computational model and our reconstituted CTF *in vitro* systems are based on human sequences. While myectomy samples from patients with R92L-cTnT and Δ160E-cTnT may be obtainable, these tissues represent very late-stage disease from patients who have likely undergone medical treatment for years and thus, would not inform on our central question regarding the earliest stages of diastolic function *in vivo*. A second approach using isogenic human derived iPSC-CMs would directly address the early-onset stage. However, one of the enduring limitations in the field remains the inability to derive myocytes with greater than 50% cTnI, and thus 50:50 ssTnI:cTnI at best. ssTnI is missing the first 30 amino acids of the N-terminus, including cTnI-S_23_S_24_, vastly complicating the functional and structural interpretation of the effects of mutations. We also note that while cTnC-T53C-IAANS is environmentally sensitive, it is not a direct measure of Ca^2+^ dissociation. Rather, it measures fluorescence intensity changes associated with an array of structural changes to cTnC during Ca^2+^ dissociation. These measurements are reflective of the milieu of structural changes induced by mutations and other alterations to the CTF. We are currently exploring the application of advanced sampling techniques in our computational approaches to directly assess the potential role of structural changes in Site II (EF-Hand motif) to address these questions.

## ACKNOWLEDGEMENTS

None.

