## Supplemental Material for "The HCM – Linked Mutation Arg92Leu in TNNT2 Allosterically Alters the cTnC – cTnI Interface and Disrupts the PKA-mediated Regulation of Myofilament Relaxation"

### SUPPLEMENTARY MATERIALS

*Experimental Animals*

All animal studies were maintained in compliance with the Public Health Service animal welfare policy, the American Association for the Accreditation of Laboratory Animal care and the Institute for Animal Care and Use. The animal research protocol was approved by the Albert Einstein College of Medicine Institute for Animal Studies and the University of Arizona University Animal Care. The C57BL/6J (RRID:IMSR_JAX:000664, Jackson Laboratories**)** background was used in all animal studies, a total of 202 mice were used. For all terminal studies, mice were anesthetized with inhaled isoflurane (3%) and respiratory rate and loss of response to stimulation were monitored to ensure appropriate anesthetic depth.

*Generation of Transgenic Mouse Models*

The missense mutation Arg92Leu (R92L) and the in-frame deletion of a glutamate residue Δ160Glu (Δ160E) located in the α-TM binding domain of cTnT were used as previously described [1,2]. Briefly, R92L-cTnT and Δ160E-cTnT express 50%, and 70% cTnT replacement, respectively and exhibit no change in the stoichiometry of cTnT. All mice were bred in-house on a congenic C57BL/6 background and transgenic mice were identified via PCR analysis. Equal numbers of male and female animals were used initially to assess the effect of sex as a biological variable on each transgenic background. Echocardiography revealed no functional change (assessed by %FS and E/e') was observed for either mutation associated with sex, males and females were pooled in subsequent studies. The cTnI-phosphomimetic transgenic mouse line (also on a C57BL/6 background) was acquired from the laboratory of Dr. Anne M. Murphy from the Johns Hopkins School of Medicine and has 95% of the cTnI residues serine-23 and serine-24 changed to aspartate-23 and aspartate-24 (cTnI-$D_{23}D_{24}$, NTDD) [3]. NTDD mice were crossed with our R92L-cTnT and Δ160E-cTnT mice to generate the double transgenic R92L-cTnT/cTnI-$D_{23}D_{24}$ (RLDD) and Δ160E-cTnT/cTnI-$D_{23}D_{24}$ (ΔDD) mice. All genotypes were viable and generated at expected Mendelian ratios. The animal research protocol was approved by the Albert Einstein College of Medicine Institute for Animal Studies and the University of Arizona University Animal Care. Animals were maintained in compliance with the Public Health Service animal welfare policy, the American Association for the Accreditation of Laboratory Animal care and the Institute for Animal Care and Use.

*Echocardiography*

To assess systolic and diastolic cardiac function mice were pre-anesthetized after hair removal (Nair) using 2-4% isoflurane prior to data collection to ensure deep anesthesia and maintained at ~1.5% isoflurane during acquisition. After anesthesia mice were placed in the prone position on a temperature regulated platform (40°C) which monitors respiratory rate and ECG and restrained using surgical tape and electrode gel (SignaGel, Parker Labs). Aquasonic (Parker Labs) ultrasound gel was applied to the chest and images were acquired in two-dimensional short-axis (SAX), targeted M-mode (in the SAX orientation) at the mid-papillary muscle, and apical four chamber orientation using a Vevo 2100 high-resolution ultra-sound imaging system (Visual Sonics, Toronto, CA) and an MS550D scan head with a tunable broadband scanning frequency range (up to 55 MHz). Using isoflurane, we regulated heart rate to separate the E and A waves for diastolic assessment. For all measurements an average of three measurements from consecutive cardiac cycles were analyzed as suggested by the American Society of Echocardiography.

*Western Blotting*

The protocol for Langendorff perfusion was adapted from Wolska et al. and Haim et al. [4,5]. Briefly, mice were heparinized (200U, Medline) fifteen minutes prior to surgery and anesthetized with 3% isoflurane. Hearts were rapidly excised via midline sternotomy and rinsed with a 4°C, pH 7.4, $Ca^{2+}$/$Mg^{2+}$-free phosphate buffered saline (PBS; Invitrogen). The blood vessels, pericardium, and thymus tissue were trimmed away in order to isolate the aorta. The aorta was cannulated with a 23-gauge leur stub adapter (Intramedic) and secured in place using 4-0 black silk suture (Deknatel). Also attached to the cannula was a 10 ml syringe (BD) containing 37°C, pH 7.4, $Ca^{2+}$ Tyrode solution (in mmol/L: NaCl 137, KCl 5.4, $CaCl_2$ 1.8, $MgCl_2$ 0.5, HEPES 10, Glucose 10). After briefly flushing the heart, the cannula was transferred to a water jacketed glass heating coil (CGS Glassware), a water bath circulator (HaakeC10, Thermo-Fisher) was used to warm up the perfusate to 37˚C and an aerator supplied 95% O2/ 5% CO2. Untreated hearts were perfused for 30 minutes, at 3 mL/min constant flow using a peristaltic pump, in $Ca^{2+}$ Tryode solution. Hearts treated with Isoproterenol, a non-selective β-adrenergic receptor agonist, were perfused with $Ca^{2+}$ Tyrode solution for 25 minutes followed by a 5-minute perfusion with $Ca^{2+}$ Tyrode solution containing 100 nmol isoproterenol. Upon conclusion of treatment atrial tissue was trimmed and ventricles were immediately flash frozen with LN2 pre-frozen Wallenberg tongs. Ventricles were immediately homogenized and sonicated in homogenization buffer (in mmol/L: Imidazole 10, Sucrose 300, NaF 25, DTT and 1 mmol/L). A protease inhibitor cocktail (12uL/mL, Thermo Scientific 78438) was added to prevent degradation of tissue homogenates. Concentrations of homogenates were determined by the Pierce BCA Assay Kit (Thermo Scientific) using bovine serum albumin as a standard. Ventricular homogenates were separated on 8-16% SDS-polyacrylamide under reducing (5%-2-Mercaptoethanol conditions, PLB and p-PLB) or nonreducing (heat treated, cTnI and p-cTnI) conditions. Proteins were transferred to a 0.2-µm nitrocellulose membrane (Thermo Scientific), stained and imaged for total protein (Revert, LI-COR, NE, USA) and incubated with primary antibody diluted in a TBS blocking buffer (LI-COR; Phospholamban, PLB, 1:1000, Thermo Fisher PA5-78410; p-Ser16 PLB, 1:1000, EMD Millipore-07-052; 1:1000, cTnI, Cell Signaling 4002S; p-Ser23/24 cTnI, 1:1000 Cell Signaling 4004S). Membranes were incubated with fluorescent secondary antibodies (1:20,000 LICOR) and visualized with the Odyssey CLx Imaging system (LI-COR, NE, USA). We then assessed the changes in expression by calculating the log fold change of the normalized (to total protein) ratios of pTnI/TnI and pPLB/PLB and the raw expression of each individual protein (TnI, pTnI, PLB, and pPLB) relative to each Non-Tg control.

*Ex Vivo Hemodynamics*

The protocol for the isolation of ventricular cardiac myocytes from 4-6 month old male and female mice was performed as above with modifications. Briefly, after cannulation of the aorta, a 25˚C, pH 7.46, $Ca^{2+}$-free Tyrode solution (in mmol/L: NaCl 113, KCl 4.7, $NaHPO_4$ dibasic 0.6, $KH_2PO_4$ monobasic 0.6, $MgSO_4$ 1.2, phenol red 0.032, $NaHCO_3$ 12, $KHCO_3$ 10, taurine 30, Hepes 10, glucose 5.55, 2,3-butanedione monoxime 9.89; Sigma-Aldrich) was used to flush the heart. The cannula was then transferred to a water jacketed glass heating coil (CGS Glassware). The heart was perfused at 3 ml/min (Rabbit Peristaltic Pump, Rainin) with a 37˚C, pH 7.4 modified Krebs-Henseliet solution (in mmol/L: NaCl 137, KCl 4, NaHCO3 24.9, MgSO4• 7H2O 1.2, glucose 11.2, CaCl2 1.8) that was aerated with 95% O2/ 5% CO2. Once perfusion of the heart had commenced, an incision was made in the left atria. To drain Thebesian flow in the left ventricular cavity, a one-inch piece of PE-50 tubing with one end partially melted was passed through the mitral valve to pierce the apex of the heart. A custom-made balloon made out of polyvinyl chloride film was also passed through the mitral valve and inflated in the left ventricle to set an end-diastolic pressure of ~10 mm Hg and held constant for the duration of the recording. The balloon was made by tying it to one end of a 12-inch piece of PE-50 tubing which had a 23-gauge cannula attached to the other end. The balloon was filled with degassed sterile water and connected to a clip-on dome with a silicone base that sits directly above a pressure transducer that was connected to the Powerlab 8/30 acquisition system (ADInstruments). A 100 µl gas tight Hamilton syringe was attached to the balloon-dome setup and was used to inflate and deflate the balloon. The balloon-dome setup was calibrated to atmospheric pressure per manufacturer instructions at the beginning of each experiment. Platinum wires (ADInstruments) were placed on the right atrium to electrically pace the heart at 400 bpm. As isolated hearts lack neurohormonal input, the cardiac functional response is insulated from external confounding effects. The heart was then submerged in a bath containing perfusion solution to maintain the temperature at 37˚C for the duration of the experiment. After 20 minutes of perfusion to allow the heart to equilibrate to its environment, increasing bolus doses (0.01, 0.1, 1.0, 10, 100 µmol/L) of dobutamine (Bedford Laboratories), a $\beta_1$-adrenergic receptor agonist, were administered every 5 min to the heart via an in-line injection port (Radnoti). Pressure development over time was acquired using the Powerlab ChartPro version 7.0 software system (ADInstruments) at a sampling rate of 1 kHz. The Blood Pressure module version 1.3 was used to compute the derivative of the pressure signal from the average of 10 consecutive tracings to compute peak positive change in pressure over time (+dP/dt) and peak negative change in pressure over time (-dP/dt).

*Protein Expression and Purification*

cDNA sequences encoding human cTnT (hcTnT), human cTnI (hcTnI), and human cTnC (hcTnC) were inserted into pET3D vectors and provided by J.D. Potter (University of Miami). Ala–Ser α-Tm wa inserted into a pET3D vector and provided by David Wieczorek (University of Cincinnati). Single cysteine substitutions were introduced to hcTnI at site A17C and A28C via the QuickChange II XL site-directed mutagenesis kit (Agilent Technologies). The cTnC(84C) construct was made via substitution of the endogenous cysteine at site 35 with a serine. Each clone was sequenced by the University of Arizona Genetics Core through direct DNA sequencing and verified using the SnapGene Viewer (GSL Biotech LLC). The cTnI, cTnC, and Ala-Ser $\alpha$-tropomyosin plasmids were transformed in to BL21 competent cells. While the cTnT plasmid was transformed into Rosetta (DE3) competent cells Novagen (EMD, Millipore). All transformed cells were streaked onto Luria Broth ampicillin agar plates and incubated at 37°C overnight. A single colony from each plate was inoculated into 5-7mL of LB media and incubated at 37°C while shaking at 250 rpm for 7 hours. 1mL of the starter culture was then inoculated into either ZYP medium (1% tryptone, 0.5% yeast, 0.5% (w/v) glycerol, 0.05% glucose, and 0.2% lactose) with 5% 20X P-buffer (1 M $Na_2HPO_4$, 1 M $KH_2PO_4$, and 0.5 M $(NH_4)_2SO_4$), 1 mM $MgSO_4$, and ampicillin (cTnT, cTnI, and Tm) or overnight Express TB medium (cTnC). The large cultures were grown overnight at 37°C shaking at 250 rpm. The large cultures were then collected and centrifuged at 4000 rpm for 20 min at 4°C.

For all purification protocols the Bio-rad LP purification system was used. The cTnI bacterial pellets were resuspended in 50mL Sp-Sepharose buffer (6M Urea, 50mM Tris, 2mM EDTA, and 1mM DTT, pH 7.0). The pellets were then frozen at -80°C and thawed for sonication. The suspended pellet was sonicated for 30 second bursts with 2-minute pauses for 6 cycles. The resuspended pellet was then centrifuged for 45 minutes at 17,000 RPM to pellet bacterial debris. The supernatant was then kept and loaded into an Sp-Sepharose column (Sigma; packed in a Bio-Rad Econo-Column with a 100 mL bed volume) at a rate of 1.3mL/min. The column was then washed with 5-7x column volume of Sp-Sepharose buffer and eluted via a linear gradient from 0-0.6M KCl in Sp-Sepharose buffer. Fractions containing cTnI were determined through Coomassie staining of sodium dodecyl sulfate polyacrylamide gel electrophoresis (SDS-PAGE) gel, which were then pooled and dialyzed against 2L TnC affinity buffer (50mM Tris, 2mM CaCl2, 0.5 M KCL 1mM DTT, pH 7.5) for two subsequent dialysis changes for at least 8 hours. Dialyzed protein was then loaded into the TnC affinity column prepared as per the manufacturer's protocol for conjugating proteins to a cyanogen bromide-activated Sepharose 4B gel (Sigma) and packed in an Econo-Column (Bio-Rad). The column was washed with 5–7× column volumes of TnC affinity buffer, and the protein was eluted using both urea and EDTA gradients (0–6 M and 0–3 mM, respectively) in TnC affinity buffer.

cTnT was purified on an Sp-Sepharose column and eluted with a linear gradient of 0–0.6 M KCl as described for cTnI. The fractions containing cTnT were determined though Coomassie staining of SDS-PAGE gel. The fractions with cTnT protein was dialyzed against 2L of Q-Sepharose buffer (6M Urea, 20mM Tris, 1mM EDTA, 1mM DTT, pH 7.8) for two subsequent dialysis changes for at least 8 hours. The dialyzed protein was recovered and loaded on a Q-Sepharose column (Sigma; packed in a Bio-Rad Econo-Column with a 100 mL bed volume). The column was then washed with 5-7x the column volume of Q-Sepharose buffer and eluted with a linear gradient of 0-0.6M KCl in Q-Sepharose buffer.

cTnC was initially purified on a Q-Sepharose column as described with cTnT. The fractions containing cTnT were determined though Coomassie staining of SDS-PAGE gel. The fractions with cTnT protein was dialyzed against 4L of Phenyl-Sepharose A buffer (50mM Tris, 1mM CaCl2, 1mM MgCl2, 50mM NaCl, 1mM DTT, pH 7.5) for four subsequent dialysis changes for at least 8 hours at 4C. During the dialysis process the room temperature phenyl Sepharose column was regenerated with 5x the volume of de-gassed 30% isopropanol (in dd$H_2O$) followed by 500mL of ddH2O. The phenyl Sepharose column was then pre-equilibrated with room temperature, degassed Phenyl Sepharose A buffer with the addition of 0.5M ammonium sulfate. The protein was recovered from dialysis and allowed to warm to room temperature. Once the protein was at room temperature, solid ammonium sulfate was added to a concentration of 0.5M. The protein was then loaded into the Phenyl-Sepharose column (Sigma; packed in a Bio-Rad Econo-Column with a 100 mL bed volume) at a rate of 1.3 mL/min. The column was then washed with 5x the column volume of the Phenyl-Sepharose A buffer with 0.5M ammonium sulfate. The cTnC was then eluted with 500mL of Phenyl Sepharose C buffer (50mM Tris, 1mM EDTA, 1mM DTT, pH 7.5) at a rate of 1.3mL/min.

Ala-ser $\alpha$ -Tm bacterial pellets were resuspended in 40 mL of dd$H_2O$ and transferred to a plastic beaker. 4 mg of lysozyme was added to the resuspended pellet and stirred on ice every 5-10minutes for an hour. The pellet was then frozen at -80°C and then thawed. Solid NaCl was added to the thawed pellet to a final concentration of 1M and then sonicated on ice for 3 minutes with 3 minutes of rest for 3 cycles. The sonicated pellet was then centrifuged at 17000 RPM for 45 minutes at 4°C to remove bacterial debris. The supernatant was then collected in a 50mL conical and boiled for 45 minutes. The sample was then centrifuged for 10 minutes at 17000 RPM at 4°C. The supernatant was collected and 1M HCL was added dropwise to a pH of 4.4-4.6 to precipitate the tropomyosin. The sample was then centrifuged for 10 minutes at 17000 RPM at 4°C. The supernatant was decanted, and the pellet was resuspended in 1M KCl. KOH was added dropwise to a pH of 7-8 to resuspend the pellet. This process was repeated 3 times in order to obtain purified tropomyosin. F-Actin was isolated and purified from rabbit skeletal muscle as previously described [6]. For all proteins purity was determined through Coomassie staining of the SDS-PAGE gels.

*Protein Labeling*

For all protein labeling initial protein concentrations were determined via UV-vis spectrophotometry, all labeling procedures were performed in the dark. For TR-FRET experiments the cys-substituted proteins hcTnI-A17C and hcTnI-A28C were labeled with acceptor probe DABMI (Setareh Biotech) and TnC(84C) was labeled with donor probe IAEDANS (Molecular Probes). The individual subunits were dialyzed into 6M UREA, 0.3M KCl 50mM MOPS 1mM EDTA at pH 7 for DABMI labeling reactions and pH 8 for IAEDANS labeling reactions. The proteins were recovered, and the concentration was determined. The reducing agent TCEP (Sigma) was then used to spike the samples at 4x the concentration of the protein and incubated at room temperature for 30 minutes. A 10x molar excess of dye was added to the protein samples and the reaction was allowed to proceed at room temperature for 2 hours. The reaction was stopped via the addition of 5mM DTT. Excess dye was dialyzed out with a minimum of three 1L dialysis changes. Excess aggregated dye was removed by spinning at 15000rpm for 15 minutes via tabletop centrifuge. Label concentration was determined via concentration of dye/initial concentration of protein. All experiments were performed with labeled troponin subunits with over 90% labeling.

IAANS labeling (Toronto Research Chemicals) of cTnC-T53C for stopped flow kinetics experiments was performed similarly with slight modifications. Briefly, cTnC-T53C was reduced via mixing with 5mM DTT for 8 hours at room temperature. The reduced sample was then dialyzed against 2L of 6M Urea, 90mM KCl, 50mM Tris base, 1mM EGTA, pH 7.5 at 4°C prior to labeling. Fivefold molar excess IAANS dye dissolved in DMF was then added dropwise to cTnC-T53C. Labeling was carried out for 5 hours with gentle mixing at room temperature. The reaction was stopped via the addition of 5mM DTT, and unreacted IAANS was removed via exhaustive dialysis against Post-Label Buffer (90mM KCl, 10mM MOPS, pH 7.0).

*Thin Filament Reconstitution*

The troponin subunits were dialyzed into reconstitution solution (6M urea, 0.5M KCl, 30mM MOPS, 3mM $MgCl_2$, 1mM DTT, pH 7.0) and then reconstituted in the ratio 1.2 TnT:1 TnC :1.2 TnI for donor-only troponins and stopped-flow troponins and 1.2 TnT:1 TnC: 1 TnI for donor-acceptor troponins. The combined troponin subunits were then put back into reconstitution solution and the urea concentration was progressively decreased from 6M to 4M to 2M to 0M with each dialysis occurring for at least 6hrs. The donor-only and donor-acceptor troponins were then dialyzed into TR-FRET buffer (0.15MKCl 50mM Mops 5mM MgCl2 2mM EGTA, and 1mM DTT, pH 7.0) and the stopped-flow
troponins were dialyzed into kinetics buffer (10mM MOPS 0.15M KCl 3mM MgCl 2, , 1mM DTT, pH 7.0). Troponins were then clarified by centrifuge at 15000RPM for 5 minutes via tabletop centrifuge.

Both F-actin and Tropomyosin were dialyzed into working buffer and the thin filaments were reconstituted at rations 0.8 Tn: 1 Tm: 7 F-actin for TR-FRET and 0.88 Tn: 1 Tm: 7 F-actin for stopped flow experiments in order to ensure all troponin was reconstituted into thin filaments. F-actin was initially incubated with tropomyosin on ice for 40 minutes and troponin was subsequently added and incubated for 20 minutes. Working buffer was then added to dilutes the thin filaments to a final working concentration of 2.5 $\mu$M IAEDANS for TR-FRET and 0.5 M IAANS for stopped-flow experiments.

*Stopped-flow Kinetics*

$Ca^{2+}$ dissociation rates ($k_{off}$) were measured using an Applied Photophysics SX20 stopped-flow machine with a deadtime of 1.25ms at 15°C. For all dissociation events, sample +200μM CaCl 2 were mixed with 12mM EGTA. The cardiac thin filaments were excited at 330nm using an excitation monochromator, and emission was monitored at 510nm using an emission monochromator. Data traces, representing an average of 5-8 serial injections, were fit with a single exponential decay function to determine each observed $k_{off}$ value. We used previously published buffer conditions for stopped-flow kinetics experiments in our system [7]; however, the use of this kinetics buffer resulted in traces that could not be fit with an exponential decay function for the ΔDD condition (Figure S3). Therefore, $Ca^{2+}$ dissociation rates for the ΔDD condition were performed with a lower salt concentration (10mM MOPS, 0.09M KCl, 3mM MgCl 2).

*Time Resolved-Fluorescence Resonance Energy Transfer*

Thin filament sample volumes were then split, and half of the samples were spiked with 1mM $CaCl_2$ in for the $Ca^{2+}$ saturated and unsaturated conditions. 100 $\mu$L of the thin filaments were loaded into a corning 96 well special optic microplate (Sigma). Lifetime measurements of donor only and donor acceptor thin filaments were performed using ISS ChronosBH time correlated single photon counting system with an integrated K428 microwell plate reader. A 375 nm laser was used to excite the samples at 30% power. Sample excitation was filtered through a 420 long pass emission filter. The pulse clock frequency was set to 5MHz with analog to digital conversion (ADC) at <1% of the repetition rate. Lifetime data was collected in the time range of 0-100ns with an offset of 3ns. Lifetime data was analyzed using FargoFit created by David Thomas' group. The donor only decay was fit to a sum of exponentials (equations 1). n =3.

$$F_D(t) = \sum_{i=1}^{n} A_i e^{\left(-\frac{t}{\tau_{D_i}}\right)} \qquad \text{Eq. 1}$$

The donor acceptor traces were then fit with equation 2 . The most optimal FRET donor acceptor population was from a one gaussian distribution of FRET distances. This fitting uses the gaussian function in equation 3.

$$F_{DA}(t) = \int p(r) \sum_{i=1}^{n} A_i e^{\left(-\frac{t}{\tau_{D_i}}\right)\left[1+\left(\frac{R_{0_i}}{R}\right)^6\right]} dr \qquad \text{Eq. 2}$$

$$p(r) = \frac{1}{Z}\frac{1}{\sigma\sqrt{2\pi}} e^{-\frac{1}{2}\left(\frac{r-\bar{r}}{\sigma}\right)^2} \qquad \text{Eq. 3}$$

*Molecular Dynamics Simulations*

Computational simulations begin with a fully atomistic model of the cardiac thin filament. A previous iteration of the model compiled structural information of the troponin core, tropomyosin, and actin together, while additionally modeling unstructured segments of Tn with homology or secondary structure prediction tools [8-10]. Alterations to the model were made due to recently published cryo-EM reconstructions of reconstituted and native thin filaments as well as corrections to unphysical Tm helical pitch present in the aforementioned cryo-EM models [11-13]. This allowed for atomic models to be determined for $Ca^{2+}$ depleted and $Ca^{2+}$ saturated cardiac thin filaments. These versions of the atomic model for the cardiac thin filament, used in the present study, were previously utilized in conjunction with fluorescence studies to examine an unstructured portion of cardiac troponin T and its orientation within the cardiac thin filament [6].

Once compiled, the WT atomic model in both $Ca^{2+}$ saturation states were prepared for MD by initially solvating the protein complex in a water box with a minimum 15 Å solute-solvent distance with the TIP3 water model [14]. Counterions of K+ and Cl- are added to neutralize the system with a set concentration of 0.15 mol/L. These processes are carried out in the visual molecular dynamics program version 2.13 with the CHARMM 36 parameters [15,16]. From here, MD is carried out within the nanoscale molecular dynamics (NAMD) program version 2.14 [17]. The prepared systems were subjected to 5000 steps of minimization with the conjugate gradient method to minimize atomic clashes, followed by a heating phase where the temperature of the system is increased at a rate of 1 K/ps until a final temperature of 300 K is achieved. A final 2 ns run is performed at a constant temperature of 300 K and a constant pressure of 1 atm to stabilize the system at the desired temperature.

The mutant structures are modified from the initial WT structures and subjected to a similar equilibration scheme as for WT. For the R92L-cTnT structure, the atomic coordinates for the arginine sidechain residue are deleted and replaced with leucine, according to the CHARMM parameters. To prepare the Δ160E-cTnT structure, the atomic coordinates for the $160^{th}$ residue are deleted, and a linkage is made to bridge the $159^{th}$ residue with the $161^{st}$. This creates a nonphysical, long bond between the two residues, but its final length and adjustments to the overall secondary structure are refined through MD. These procedures for creating mutations have been utilized extensively in the past [18-20].

Once each system is fully equilibrated, the measured FRET distances are incorporated into the model as follows: distance constraints are applied to the alpha carbon atoms of the measured FRET pairs. An initial 1 ns run is performed with the FRET distance constraints in place, this may be thought of as a steered MD simulation from the initial assumed model to the actual experimentally measured FRET distances. In further simulation we hold the distance constraints at the measured FRET distances, but moderate force constants in the constraint allow the FRET pairs to fluctuate 1-2 Å. This is run for another 2 ns. From this 2 ns run, the average atomic positions for each system are calculated, minimized, and examined.

*Statistical Analysis*

All values are reported as mean ± S.E.M. and were calculated using GraphPad Prism 9.5.1 (San Diego, CA). For animal studies, sample sizes were determined using *a priori* power analysis as described in the Major Resources Table when possible. All datasets were checked for normality using a Shapiro-Wilks test, and outliers were removed using a ROUT method (Q=5% unless otherwise stated) if necessary. 2D echocardiography results were analyzed using a 2-way ANOVA with Tukey correction for multiple comparisons (MC) to assess the effect of the mutation within timepoint. For western blot studies, log fold change was calculated respective to each Non-Tg control and a 2-way ANOVA with Tukey or Sidak correction for MC was used to assess the effect of the mutation within isoproterenol treatment or the effect of isoproterenol treatment within mutation respectively. For the isovolumic studies, an average of 10 consecutive traces was used for each sample and a mixed-model RM 2-way ANOVA with Tukey or Dunnet correction for MC was used to assess the effect of genotype within a dobutamine dose or the effect of increasing dobutamine dose vs baseline within genotype, respectively. $Ca^{2+}$ dissociation experiments used a 2-way ANOVA with Sidak correction for MC was used to assess the effects of genotype and phosphomimetic. For the WT TR-FRET experiments a 2-way ANOVA with Tukey or Sidak correction for MC was used to assess changes in distance and order (FWHM) within biochemical condition and across within FRET site, respectively. Lastly, for the mutation TR-FRET studies a 2-way ANOVA with Tukey correction for MC was used to probe the effect of biochemical condition and mutation for each genotype individually.

**Figure S1:** Total cTnI, Total PLB, Phospho-troponin I (pTnI) and phospho-phospholamban (pPLB) quantitation in cardiac whole homogenates for Non-Tg, R92L-cTnT, and Δ160E-cTnT reported as fold change from each Non-Tg control. Summary of A: total TnI, B: total pTnI, C: total PLB, and D: total pPLB. Values are reported as mean ± S.E.M., n = 5 animals per group. A 2-way ANOVA with Tukey correction was used to compare each genotype to their respective Non-Tg controls within treatment group, and a Sidak correction was used to compare baseline and isoproterenol treatment within genotype. A summary of adjusted p-values can be found in Table S10. * $p<0.05$, ** $p<0.01$ vs Non-Tg within -ISO; # $p<0.05$, ## $p<0.01$, ### $p<0.005$ vs Non-Tg within +ISO; † $p<0.05$ vs -ISO within genotype.

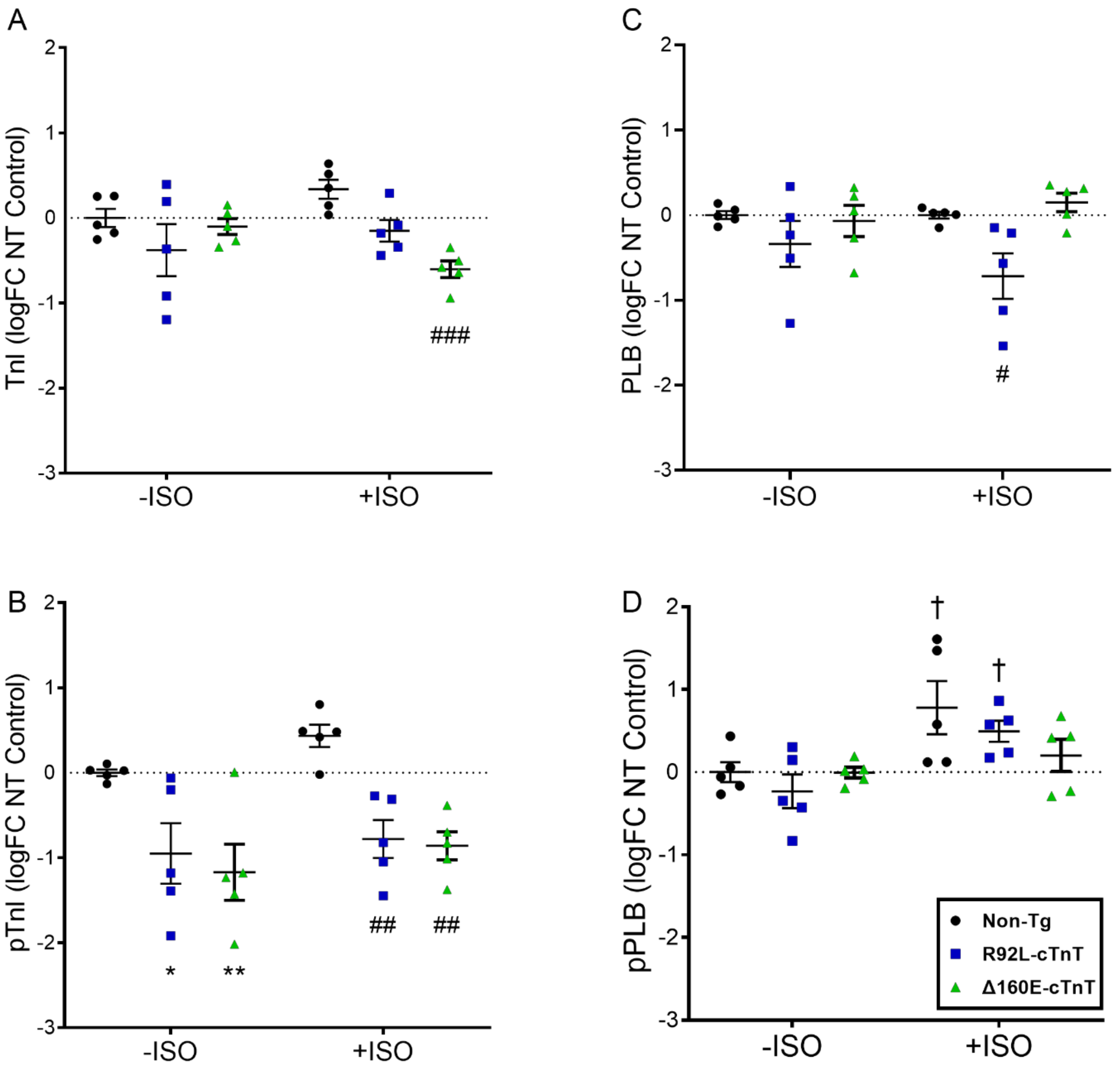

**Figure S2:** Gross morphological assessment of R92L-cTnT and Δ160E-cTnT mice and phosphomimetic crosses. A: Heart Weight/Body Weight (HW/BW) taken at approximately 4 months of age from Non-Tg, NTDD, R92L-cTnT, RLDD, Δ160E-cTnT, and ΔDD mice. B: Atria Weight/Heart Weight (AW/HW). Values are reported as mean ± S.E.M, n = 7-10 per genotype with Non-Tg littermates combined. A 1-way ANOVA with a Dunnett correction for multiple comparisons was used to compare to Non-Tg. Means and adjusted p-values are summarized in Table S11 and S12. * p<0.05, *** p < 0.001, **** p < 0.0001 vs Non-Tg.

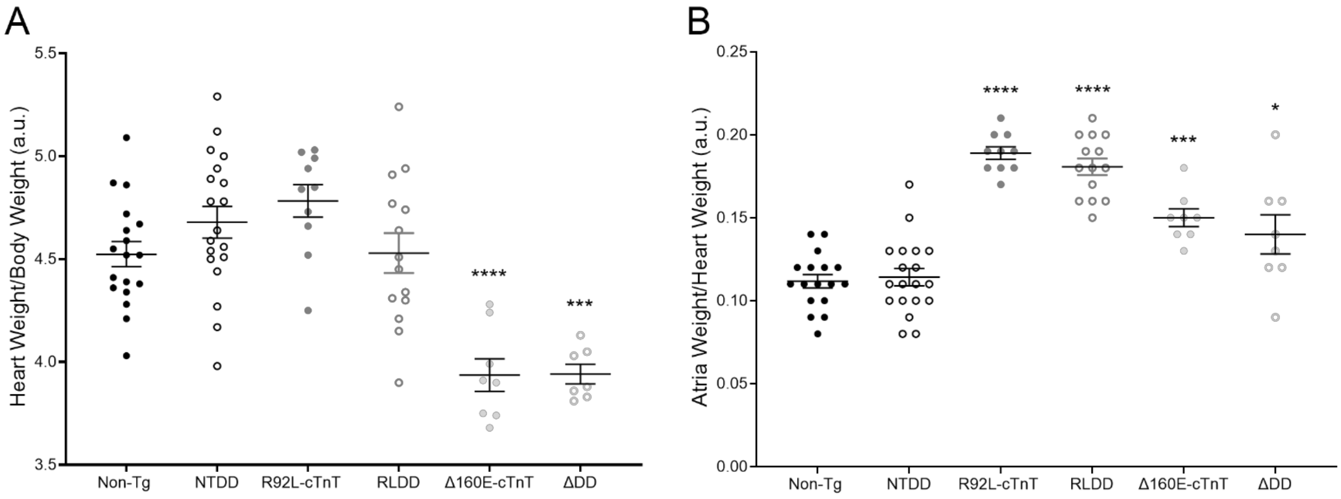

**Figure S3:** Representative traces of calcium dissociation rates from reconstituted thin filaments at low and high ionic strength demonstrating the variability in exponential decay observed under typical ionic strength conditions for A: WTDD thin filaments, B: RLDD thin filaments, and C: ΔDD thin filaments.

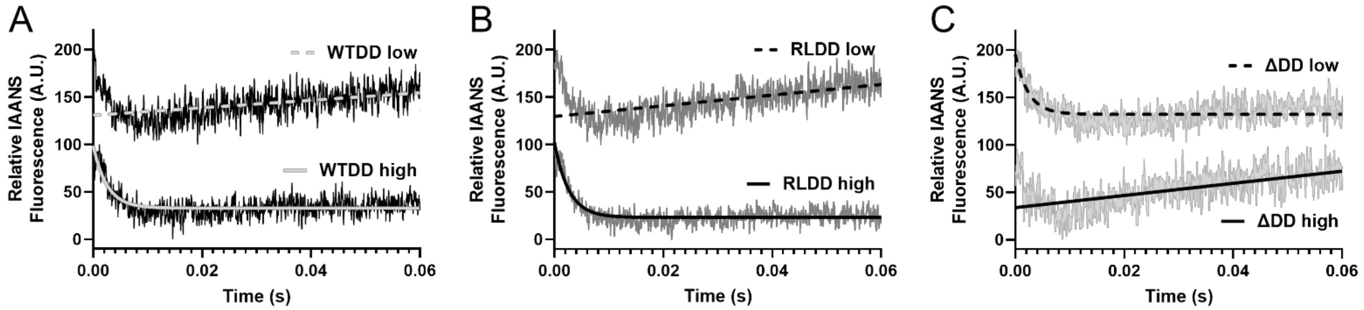

**Table S1:** Summary of echocardiographic values from Figure 1. Values are reported mean ± S.E.M, n = 5-8 per group. A 2-way ANOVA with Tukey correction was used as described above. * vs Non-Tg 1-2 months, # vs Non-Tg 4-6 months, ‖ no significance.

| | Early | | | Late | | |
|---|---|---|---|---|---|---|
| | **Non-Tg** | **R92L-cTnT** | **Δ160E-cTnT** | **Non-Tg** | **R92L-cTnT** | **Δ160E-cTnT** |
| **E/e'** | -32.7 ± 0.838 | -39.0 ± 0.991 * | -32.6 ± 0.779 ‖ | -32.1 ± 1.11 | -38.0 ± 2.02 # | -38.0 ± 1.71 # |
| **E (mm/s)** | 631.4 ± 19.8 | 553.3 ± 20.4 ‖ | 538.4 ± 30.2 ‖ | 641.8 ± 23.4 | 543.7 ± 35.7 # | 602.2 ± 27.2 ‖ |
| **e' (mm/s** | -19.4 ± 0.923 | -14.3 ± 0.797 ** | -16.6 ± 0.871 ‖ | -20.0 ± 0.443 | -14.3 ± 0.686 ### | -16.1 ± 1.31 # |
| **%FS** | 28.2 ± 0.931 | 39.2 ± 1.741 **** | 36.5 ± 0.663 **** | 25.6 ± 0.201 | 41.7 ± 0.882 #### | 38.9 ± 0.720 #### |
| **%EF** | 55.4 ± 1.419 | 70.9 ± 1.982 **** | 67.5 ± 0.884 **** | 51.0 ± 0.351 | 73.8 ± 1.103 #### | 70.6 ± 0.919 #### |
| **$T_d$ (mm)** | 0.784 ± 0.016 | 0.755 ± 0.021 ‖ | 0.773 ± 0.023 ‖ | 0.824 ± 0.024 | 0.926 ± 0.034 # | 0.826 ± 0.008 ‖ |

**Table S2:** Summary of adjusted p-values from the echocardiographic values in Figure 1 and Table S1. A 2-way ANOVA with Tukey correction was used as described above. * vs Non-Tg 1-2 months, # vs Non-Tg 4-6 months, ‖ no significance.

| | Early | | | Late | | |
|---|---|---|---|---|---|---|
| | **Non-Tg** | **R92L-cTnT** | **Δ160E-cTnT** | **Non-Tg** | **R92L-cTnT** | **Δ160E-cTnT** |
| **E/e'** | n/a | * 0.0122 | ‖ 0.9988 | n/a | # 0.0102 | # 0.0137 |
| **E (mm/s)** | n/a | ‖ 0.1846 | ‖ 0.0731 | n/a | # 0.0489 | ‖ 0.6071 |
| **e' (mm/s** | n/a | ** 0.0016 | ‖ 0.0799 | n/a | ### 0.0002 | # 0.0116 |
| **%FS** | n/a | **** <0.0001 | **** <0.0001 | n/a | #### <0.0001 | #### <0.0001 |
| **%EF** | n/a | **** <0.0001 | **** <0.0001 | n/a | #### <0.0001 | #### <0.0001 |
| **$T_d$ (mm)** | n/a | ‖ 0.6929 | ‖ 0.9473 | n/a | # 0.0121 | ‖ 0.9978 |

**Table S3:** Summary of adjusted p-values from the western blots in Figure 2. A 2-way ANOVA with Tukey or Sidak correction was used as described above. For samples with no significance (‖) the p-value referenced is vs Non-Tg -ISO. * vs Non-Tg within -ISO; # vs Non-Tg within +ISO; † vs -ISO within genotype; ‡ vs R92L-cTnT (across genotype) within +ISO.

| | pTnI/TnI | | | pPLB/PLB | | |
|---|---|---|---|---|---|---|
| | **Non-Tg** | **R92L-cTnT** | **Δ160E-cTnT** | **Non-Tg** | **R92L-cTnT** | **Δ160E-cTnT** |
| **- ISO** | n/a | ‖ 0.1159 | ** 0.0020 | n/a | ‖ 0.9308 | ‖ 0.9761 |
| **+ ISO** | ‖ 0.9797 | # 0.0372 | ‖ 0.4160 | ‖ 0.4168 | * 0.0396<br>†† 0.035 | ‡‡ 0.0021 |

**Table S4**: Summary of adjusted p-values of inotropic and lusitropic response values from Figure 3 and Table 1. A mixed-effect RM 2-way ANOVA with Dunnett's or Tukey correction was used as described above. ǁ not significant (for simplicity only significant p-values are shown), * vs baseline dobutamine dose (0 µmol/L) within mutation, # vs Non-Tg within dobutamine dose, † vs NTDD within dobutamine dose, ‡ vs R92L-cTnT within dobutamine dose, § vs Δ160E-cTnT within dobutamine dose. Where applicable for the Non-Tg and NTDD data sets, the highest (least significant) p-value from the separate ANOVAs was reported.

| | **Non-Tg** | **NTDD** | **R92L-cTnT** | **RLDD** | **Δ160E-cTnT** | **ΔDD** |
|---|---|---|---|---|---|---|
| **Dobutamine (µmol/L)** | **Maximum +dP/dt (mmHg/sec)** | | | | | |
| **0** | n/a | ǁ | ### 0.0006<br>† 0.0262 | #### <0.0001<br>†††† <0.0001 | ǁ | ǁ |
| **0.01** | *** 0.0005 | **** <0.0001 | **** <0.0001 | *** 0.0001<br>‡‡ 0.0014 | *** 0.0003 | ** 0.0019<br>† 0.0292 |
| **0.1** | *** 0.0001 | **** <0.0001 | **** <0.0001 | ** 0.0019<br>## 0.0030<br>‡‡ 0.0014 | *** 0.0009 | †† 0.0010 |
| **1** | **** <0.0001 | **** <0.0001 | *** 0.0001<br>† 0.0167 | * 0.0116<br>‡ 0.0349 | **0.0099 | ## 0.0051<br>†††† <0.0001 |
| **10** | **** <0.0001 | **** <0.0001 | **** <0.0001<br>## 0.0012<br>††† 0.0005 | **** <0.0001<br>‡‡ 0.0033 | **** <0.0001<br>## 0.0019<br>††† 0.0008 | #### <0.0001<br>†††† <0.0001<br>§ 0.0105 |
| **100** | **** <0.0001 | **** <0.0001 | **** <0.0001<br># 0.0111<br>††† 0.0004 | **** <0.0001<br>‡‡ 0.0027 | **** <0.0001<br># 0.0154<br>††† 0.0006 | * 0.0120<br>### 0.0006<br>†††† <0.0001<br>§ 0.0287 |
| **Dobutamine (µmol/L)** | **Maximum -dP/dt (mm Hg/sec)** | | | | | |
| **0** | n/a | ǁ | ǁ | † 0.0149<br>‡‡‡ 0.0006 | # 0.0186<br>†††† <0.0001 | † 0.0213<br>§§ 0.0022 |
| **0.01** | *** 0.0010 | **** <0.0001 | *** 0.0002<br>†††† <0.0001 | ** 0.0083<br>‡‡‡ 0.0006 | ## 0.0050<br>†††† <0.0001 | * 0.0479 |
| **0.1** | *** 0.0003 | *** 0.0001 | *** 0.0008<br># 0.0215<br>††† 0.0002 | * 0.0260<br>‡‡‡ 0.0003 | ## 0.0017<br>†††† <0.0001 | §§§ 0.0002 |
| **1** | *** 0.0002 | **** <0.0001 | * 0.0260<br>## 0.0022<br>†††† <0.0001 | * 0.0297<br>† 0.0121<br>‡‡‡ 0.0002 | * 0.0124<br># 0.0151<br>††† 0.0005 | †† 0.0025 |
| **10** | **** <0.0001 | **** <0.0001 | **** <0.0001<br>## 0.0037<br>†† 0.0024 | **** <0.0001<br>‡ 0.0213 | **** <0.0001<br>## 0.0021<br>††† 0.0005 | * 0.0193<br>## 0.0036<br>†† 0.0025 |
| **100** | **** <0.0001 | ****<0.0001 | **** <0.0001<br>## 0.0072<br>†† 0.0016 | **** <0.0001<br>‡‡ 0.0036 | **** <0.0001<br>## 0.0024<br>††† 0.0001 | * 0.0140<br>## 0.0043<br>††† 0.0009 |

**Table S5**: Summary of adjusted p-values from $Ca^{2+}$ dissociation values in Figure 4. A 2-way ANOVA with Sidak correction for MC was used as described above. * vs WT-cTnT within cTnI-WT; # vs WT-cTnT within cTnI-WTDD; † vs cTnI-WT within cTnT genotype.

| | **WT-cTnT** | **R92L-cTnT** | **Δ160E-cTnT** |
|---|---|---|---|
| **cTnI-WT** | n/a | * 0.0111 | * 0.0126 |
| **cTnI-WTDD** | †††† <0.0001 | ǁ 0.2599<br>†††† <0.0001 | #### <0.0001<br>†††† <0.0001 |

**Table S6:** Summary of distance and FWHM values obtained for WT CTFs via TR-FRET between TnC84C-TnIA9C, TnC84C-TnIA17C, and TnC84C-TnIA28C in Figure 5. Values are reported as mean ± S.E.M., n = 8-14 per group. A 2-way ANOVA with Tukey or Sidak correction as described above. A summary of adjusted p-values can be found in Table S5. * vs A9C within -Calcium, # vs A9C within +Calcium, and † vs -Calcium within site.

| **Distance (Å)** | **cTnC84C-cTnIA9C** | **cTnC84C-cTnIA17C** | **cTnC84C-cTnIA28C** |
|---|---|---|---|
| **- Calcium** | 52.2 ± 2.114 | 43.2 ± 0.868 **** | 41.8 ± 0.620 **** |
| **+ Calcium** | 48.3 ± 1.143 † | 39.1 ± 0.534 #### †† | 40.9 ± 0.428 #### |
| **FWHM (Å)** | **cTnC84C-cTnIA9C** | **cTnC84C-cTnIA17C** | **cTnC84C-cTnIA28C** |
| **- Calcium** | 22.2 ± 1.157 | 17.8 ± 1.113 * | 14.8 ± 1.249 **** |
| **+ Calcium** | 21.2 ± 1.026 | 14.7 ± 0.697 ### | 11.2 ± 0.601 #### |

**Table S7:** Summary of adjusted p-values from distance distributions in Figure 5 and Table S4. A 2-way ANOVA with Tukey or Sidak correction for MC was used as described above. For values with no significance (‖) the p-value reported is relative to A9C -Calcium. * vs A9C within -Calcium, # vs A9C within +Calcium, and † vs -Calcium within site.

| **Distance (Å)** | **cTnC84C-cTnIA9C** | **cTnC84C-cTnIA17C** | **cTnC84C-cTnIA28C** |
|---|---|---|---|
| **- Calcium** | n/a | **** <0.0001 | **** <0.0001 |
| **+ Calcium** | † 0.0475 | #### <0.0001<br>†† 0.0029 | #### <0.0001 |
| **FWHM (Å)** | **cTnC84C-cTnIA9C** | **cTnC84C-cTnIA17C** | **cTnC84C-cTnIA28C** |
| **- Calcium** | n/a | * 0.0150 | **** <0.0001 |
| **+ Calcium** | ‖ 0.0506 | ### 0.0003 | #### <0.0001 |

**Table S8:** Summary of distance and FWHM values obtained for R92L-cTnT and Δ160E-cTnT filaments via TR-FRET between cites TnC84C-TnIA17C and TnC84C-TnIA28C in Figure 6. Values are reported as mean ± S.E.M., n = 9-14 per group. A 2-way ANOVA with Tukey correction was used as described above. A summary of adjusted p-values can be found in Table S7. * vs WT -Calcium; # vs R92L-cTnT -Calcium; † vs Δ160E-cTnT -Calcium.

| | **cTnC84C-cTnIA17C** | | | **cTnC84C-cTnIA28C** | | |
|---|---|---|---|---|---|---|
| **Distance (Å)** | **WT** | **R92L-cTnT** | **Δ160E-cTnT** | **WT** | **R92L-cTnT** | **Δ160E-cTnT** |
| **- Calcium** | 43.2 ± 0.868 | 42.1 ± 1.123 ‖ | 42.9 ± 1.340 ‖ | 41.8 ± 0.620 | 42.5 ± 0.715 ‖ | 42.9 ± 0.859 ‖ |
| **+ Calcium** | 39.1 ± 0.534 * | 38.7 ± 1.004**# | 38.2 ± 0.965*† | 40.9 ± 0.428 ‖ | 39.7 ± 0.680 ## | 42.2 ± 1.091 ‖ |
| **FWHM (Å)** | **WT** | **R92L-cTnT** | **Δ160E-cTnT** | **WT** | **R92L-cTnT** | **Δ160E-cTnT** |
| **- Calcium** | 18.2 ± 1.303 | 18.4 ± 1.733 ‖ | 18.2 ± 1.482 ‖ | 14.7 ± 1.177 | 14.0 ± 0.805 ‖ | 15.4 ± 1.219 ‖ |
| **+ Calcium** | 14.9 ± 0.669 ‖ | 11.7 ± 1.709 *# | 14.2 ± 0.832 † | 11.2 ± 0.601 ‖ | 11.1 ± 0.965 * | 14.6 ± 0.899 ‖ |

**Table S9:** Summary of adjusted p-values from distance distributions in Figure 6 and Table S6. A 2-way ANOVA with Tukey correction was used as described above. For values with no significance (‖) the p-values reported are relative to WT -Calcium for the respective FRET pair. * vs WT -Calcium, # vs R92L-cTnT within -Calcium, † vs Δ160E-cTnT within -Calcium. Where applicable for WT values the highest (least significant) p-value is reported.

| | **cTnC84C-cTnIA17C** | | | **cTnC84C-cTnIA28C** | | |
|---|---|---|---|---|---|---|
| **Distance (Å)** | **WT** | **R92L-cTnT** | **Δ160E-cTnT** | **WT** | **R92L-cTnT** | **Δ160E-cTnT** |
| **- Calcium** | n/a | ‖ 0.8313 | ‖ 0.9953 | n/a | ‖ 0.6273 | ‖ 0.7065 |
| **+ Calcium** | * 0.0140 | ** 0.0046<br># 0.0466 | * 0.0211<br>† 0.0496 | ‖ 0.8415 | ## 0.0051 | ‖ 0.9768 |
| **FWHM (Å)** | **WT** | **R92L-cTnT** | **Δ160E-cTnT** | **WT** | **R92L-cTnT** | **Δ160E-cTnT** |
| **- Calcium** | n/a | ‖ 0.9999 | ‖ 0.9732 | n/a | ‖ 0.9887 | ‖ 0.9809 |
| **+ Calcium** | ‖ 0.3821 | * 0.0139<br># 0.0114 | † 0.0334 | ‖ 0.1078 | * 0.0299 | ‖ 0.9878 |

**Table S10**: Summary of adjusted p-values from western blots in Supplementary Figure 1. A 2-way ANOVA with Tukey or Sidak was used as described above. For samples with no significance (‖) the p-value referenced is vs Non-Tg -ISO. * vs Non-Tg within -ISO; # vs Non-Tg within +ISO; † vs -ISO within genotype.

| | **TnI** | | | **PLB** | | |
|---|---|---|---|---|---|---|
| | **Non-Tg** | **R92L-cTnT** | **Δ160E-cTnT** | **Non-Tg** | **R92L-cTnT** | **Δ160E-cTnT** |
| **- ISO** | n/a | ‖ 0.2013 | ‖ 0.8836 | n/a | ‖ 0.3202 | ‖ 0.9471 |
| **+ ISO** | ‖ 0.3793 | ‖ 0.0786 | ### 0.0007 | ‖ >0.9999 | # 0.0176 | ‖ 0.7801 |
| | **pTnI** | | | **pPLB** | | |
| **- ISO** | n/a | * 0.0172 | ** 0.0035 | n/a | ‖ 0.6039 | ‖ 0.9996 |
| **+ ISO** | ‖ 0.4934 | ## 0.0025 | ## 0.0014 | * 0.0240 | ‖ 0.4777<br>† 0.0376 | ‖ 0.0767 |

**Table S11**: Summary of gross morphological assessment of heart weight and atrial mass from Figure S2. Values are reported mean ± S.E.M., n = 7-19 per group. Statistics are the result of a 1-way ANOVA with Sidak correction for MC. ‖ not significant vs any group; * vs Non-Tg.

| | **Non-Tg** | **NTDD** | **R92L-cTnT** | **RLDD** | **Δ160E-cTnT** | **ΔDD** |
|---|---|---|---|---|---|---|
| **HW/BW** | 4.52 ± 0.061 | 4.68 ± 0.077 ‖ | 4.78 ± 0.079 ‖ | 4.53 ± 0.097 ‖ | 3.94 ± 0.080 **** | 3.94±0.048 *** |
| **AW/HW** | 0.112±0.004 | 0.114±0.005 ‖ | 0.189±0.004 **** | 0.181±0.005 **** | 0.150±0.005 *** | 0.140±0.012 * |

**Table S12**: Summary of adjusted p-values from morphology in Figure S2 and Table S9. ‖ not significant vs any group; * vs Non-Tg.

| | **Non-Tg** | **NTDD** | **R92L-cTnT** | **RLDD** | **Δ160E-cTnT** | **ΔDD** |
|---|---|---|---|---|---|---|
| **HW/BW** | n/a | ‖ 0.5556 | ‖ 0.1741 | ‖ >0.9999 | **** <0.0001 | *** 0.0002 |
| **AW/HW** | n/a | ‖ >0.9999 | **** <0.0001 | **** <0.0001 | *** 0.0003 | * 0.0127 |
